# Toehold-Mediated Strand Displacement in Random Sequence Pools

**DOI:** 10.1101/2022.10.22.513323

**Authors:** Thomas Mayer, Lukas Oesinghaus, Friedrich C. Simmel

**Author notes:** **Corresponding Author** Friedrich C. Simmel, School of Natural Sciences, Department of Bioscience, Technical University Munich, 85748 Garching, Germany. DNA nanotechnology – Chemical Reaction Networks.

## Abstract

Toehold-mediated strand displacement (TMSD) has been used extensively for molecular sensing and computing in DNA-based molecular circuits. As these circuits grow in complexity, sequence similarity between components can lead to cross-talk causing leak, altered kinetics, or even circuit failure. For small non-biological circuits, such unwanted interactions can be designed against. In environments containing a huge number of sequences, taking all possible interactions into account becomes infeasible. Therefore, a general understanding of the impact of sequence backgrounds on TMSD reactions is of great interest. Here, we investigate the impact of random DNA sequences on TMSD circuits. We begin by studying individual interfering strands and use the obtained data to build machine learning models that estimate kinetics. We then investigate the influence of pools of random strands and find that the kinetics are determined by only a small subpopulation of strongly interacting strands. Consequently, their behavior can be mimicked by a small collection of such strands. The equilibration of the circuit with the background sequences strongly influences this behavior, leading to up to one order of magnitude difference in reaction speed. Finally, we compare two established and a novel technique that speed up TMSD reactions in random sequence pools: a three-letter alphabet, protection of toeholds by intramolecular secondary structure, or by an additional blocking strand. While all of these techniques were useful, only the latter can be used without sequence constraints. We expect that our insights will be useful for the construction of TMSD circuits that are robust to molecular noise.

## Introduction

Toehold-mediated strand displacement (TMSD) is the process by which a single-stranded (ss) nucleic acid invader strand binds to an overhang – called toehold – of a substrate strand in a double-stranded (ds) complex and displaces an incumbent strand via a branch migration process. This results in a complex of the invader and substrate and an unbound incumbent (Figure 1a). This type of reaction, though simple in principle, has been used in the implementation of a wide variety of in vitro molecular computing applications such as logic gates^1–3^, robotics^4^, neural networks5^−8^ and other dynamic and complex functions.^9–12^ The larger such systems grow, the higher the chance that different circuit components show unwanted interactions with each other. In an in vitro context, all components are typically highly purified and have freely choosable sequences, and thus such interactions can be explicitly designed against.^13^ Alternatively, circuits can be designed to be generally robust with respect to an unknown sequence background. Systems developed by Seelig et al. and Zhang et al. have, for example, been shown to remain functional in the presence of a background of total cellular RNA.^3,14^ TMSD systems have also been used extensively for in vitro sensing applications and, in the last decade, increasingly for in cellulo computing and gene regulation.^15–19^ For such applications, the presence of a large number of interfering background sequences is an unavoidable feature of the operating environment and their impact on strand displacement kinetics are difficult to anticipate. A better understanding of how a background of random sequences influences TMSD reactions and how one might design systems to be robust against such a background is therefore of great interest. For systems using only purified components, numerous studies have analyzed the kinetics of TMSD reactions for DNA^20–23^, RNA^24,25^ and RNA:DNA hybrids^25^ and also complexes containing mismatches.^26,27^ The software package NUPACK^28,29^ is widely used for equilibrium state analysis and the design of strand displacement circuits. By itself, however, NUPACK does not predict kinetics. For this purpose, tools like Multistrand^30^, KinDA^31^, and OxDNA^32^ have been used, and the factors determining the kinetics of strand displacement systems in clean environments are well understood. There is much less work on the behavior of systems containing a pool of background sequences. Such pools may occur in the context of a variety of applications, for instance when operating large in vitro DNA or RNA computing networks ^5,6,33^, in DNA storage^34,35^, biosensing applications^36^, and when performing strand displacement reactions inside living cells^16,18^. Previous studies have focused on distinguishing closely related input sequences^37,38^ using special design paradigms and simulations, such as using clamps to reduce leak and mismatch strategies to fine-tune displacement kinetics.^1,10,13,39^ However, mismatch strategies can reduce specificity and clamps do not enhance kinetics. For circuits sensing naturally occurring RNAs in cellulo, where either the circuit components or the sensed RNA can be degraded in minutes depending on the cell and RNA type^40^, defined kinetics are necessary for reliable results. The kinetics of in vitro TMSD-based sensing circuits that operate in total RNA are altered by the presence of non-target nucleic acids belonging to classes such as mRNA, rRNA, long-noncoding RNA, and so forth. Pure *in vitro* circuits have lately been used for performing neural network-like computation and such networks can theoretically be of arbitrary complexity with hundreds or even thousands of strands and complexes involved.^6^ Therefore, a strict orthogonal design is often not feasible even for well-defined *in vitro* circuits and some compromises have to be made. In this work, we will study undesired interactions of TMSD circuit components with a background of nucleic acids, try to gain a qualitative understanding of their features, and provide techniques for how to minimize them. In principle, one could consider explicitly analyzing the interactions of circuit components with an actual transcriptome, e.g., that of HEK293 cells, or within one particular large strand displacement circuit. Instead, we here abstract these interactions by using random pools of single-stranded DNA of a known length and concentration.

**Figure 1.**
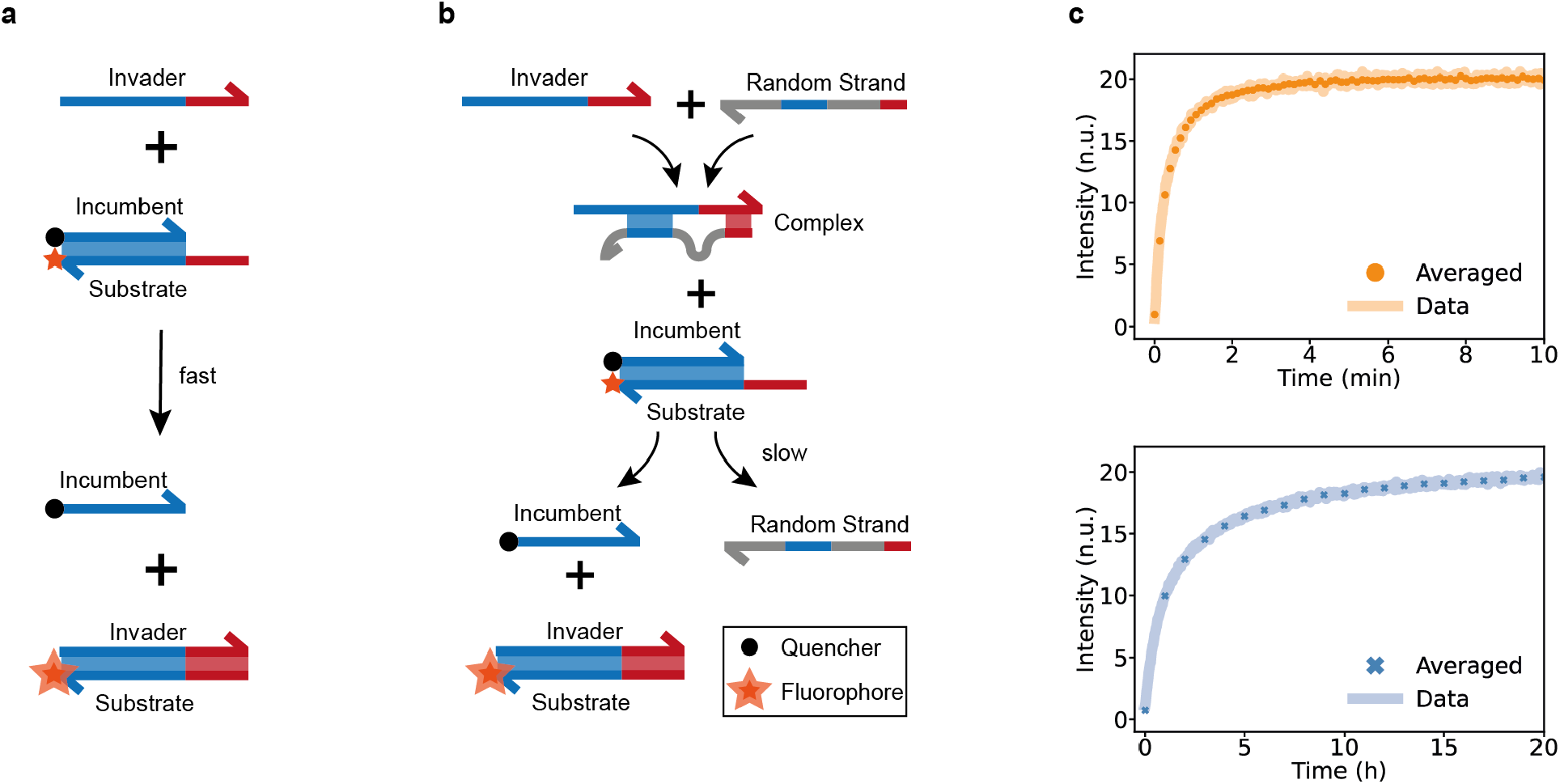
Toehold mediated strand displacement (TMSD) with and without background sequences. (a) Schematic of a TMSD process in a clean environment. The toehold of the single-stranded (ss) invader (red) binds to the complementary part of the substrate which is in a double-stranded (ds) complex with the incumbent. After initial binding of the toehold, the branch migration domains (blue) undergo a random walk-like displacement process. Eventually the invader fully displaces the incumbent due to the higher thermodynamic stability caused by the toehold, separating the quencher from the fluorophore, leading to an increase in fluorescent signal. (b) Schematic of a TMSD process in a complex environment containing background sequences. The ss invader binds to an ssDNA of the random pool, forming a partially ds complex. The newly formed complex undergoes the displacement process with a less accessible toehold and a partially bound branch migration domain. (c) Fluorescence intensity (F.I.) measurements of a TMSD system without a random sequence pool (top) and after first equilibrating the invader with a random sequence pool (bottom) in normalized units (n.u.). We used an invader concentration of 20 nM and a reporter (substrate + invader) concentration of 40 nM for all kinetic measurements. The normalized units show the concentration of invader that has already reacted. Note that the time scale for the reaction is slowed down from minutes to hours by the random pool. All measurements were performed at 29°C.

This abstraction has two motivations: First, in considering a specific background, we would lose generality as the sequence backgrounds even in purified total RNA, in the cytoplasm, or the nucleoplasm are very different even for a single cell type^41^ and the same is true for different TMSD circuits. We are thus interested in the more general impact of random sequences across a wide range of possible environments and applications. Second, focusing on random sequence pools simplifies both the computational and the experimental analysis, allowing for a deeper understanding of the underlying processes. This abstraction approach has previously been used to test the robustness of reaction networks with respect to non-specifically interfering sequences, but has not been studied in greater detail.^42,43^ **Figure 1**c gives an impression of the impact of a high concentration of random ssDNA background strands of 50 nt length on the kinetics of a one-step TMSD circuit when the invader and the background were pre-equilibrated. The bare circuit (i.e., in absence of a random pool) runs to completion in a few minutes. In the presence of the background, random strands bind to circuit components, slowing down the reaction (**Figure 1**b). This is caused by the less accessible toehold and the branch migration domain partially undergoing 4-way instead of the usual 3-way branch migration (Figure S1). The invader strand is particularly vulnerable because it is fully single-stranded by design. Consequently, the circuit takes almost one day to run to completion, corresponding to a slowdown by well over a factor of 100.

In the present work, we investigate the details of the underlying processes using various approaches. We begin by studying the impact on TMSD kinetics of individual, NUPACK-designed interfering strands that interact with the invader to form a defined secondary structure. From the experiments, we heuristically derive features that classify the interaction and use them as inputs for different predictive models (regression trees, symbolic regression, and neural networks). By evaluating the performance of these models, we show that a small collection of understandable features can be used to gain intuition about kinetics. Next, we investigate how different lengths and concentrations of random DNA influence the kinetics. The results differ strongly depending on whether or not the invader strand is pre-equilibrated with the random strands. For the equilibrated case, we hypothesize, based on NUPACK simulations, that only a small subset of all random strands is relevant to kinetics. We further illustrate this by designing representative sets of strands that recapitulate the energy landscape of the random strands in the experiment, and indeed have a very similar impact on kinetics as the random pool. Based on our insights, we finally study three different approaches that successfully render TMSD systems resilient against background sequences.

## Results

### Impact of individual interfering strands on reaction kinetics

The interaction between the TMSD circuit components and the random sequence pool is multifaceted, with the most important aspect being the interaction between the invader and the background nucleic acids. As the invader is commonly designed to be a single strand free of secondary structure, it is particularly vulnerable to binding interactions. While it is not necessary to use toeholds longer than eight bases for regular in vitro strand displacement circuits (the reaction does not speed up for longer toeholds), in cellulo applications often use up to 20 bases in the toehold.^18,44^ We therefore chose to conduct our experiments using rather long toeholds of 10 or 20 bases. We also decided to concentrate on the interaction of the invader strand with the strands of the random pool as the dominant interaction, as the reporter complex - consisting of substrate and incumbent - does not show strong binding to the molecular background (Figure S2). Further, as the reporter is present in excess relative to the invader, the kinetics would not be altered even if a fraction of the reporter toeholds was sequestered by random strands.

To experimentally study the effect of individual strands taken from a random sequence background on reaction kinetics, we used NUPACK to design ≈200 strands with a length of 35 nt that form a variety of different secondary structures with the invader strand, and ordered them from an oligo supplier. We individually incubated each of the strands with the invader strand and then added the reporter to produce kinetic curves via fluorescence measurements. We fitted our data as a one-step second order reaction with an effective kinetic rate constant (k_eff_). Although the reactions are expected to be more complex in detail, assigning a single effective kinetic constant greatly simplified downstream analysis and turned out to be a sufficient measure for kinetics in the context of our study. We repeated the same experiment for a second strand displacement circuit using a different set of ≈100 interfering strands.

The kinetic data obtained from these ≈ 300 designed strands represent a mapping between secondary structure and reaction speed. Using many more measurements for a much larger number of circuits than this would ultimately allow us to train an end-to-end machine learning model that predicts kinetics directly based on the secondary structure or even the interfering strand sequence. A priori it is not clear, however, whether such a model would accurately predict the kinetics of drastically different circuit designs and environments. Thus, we were here interested in developing an intuitive understanding of the impact of interfering strands on kinetics based on explainable structural features.

A few examples with different secondary structures and their influence on displacement kinetics can be seen in **Figure 2**a. The primary insight obtained from these three examples is that, as the number of bound toehold bases decreases, reaction speed increases despite additional bases bound in the branch migration domain, indicating that secondary structure in the toehold domain has a higher impact on kinetics. This is not unexpected since the dependence of the reaction speed of “bare” strand displacement circuits (in the absence of background) is exponential for short toehold lengths.^45^ Secondary structure in the branch migration domain results in a combination of three- and four-way branch migration (3WBM and 4WBM) processes during the displacement of the incumbent (visualized in Figure S1). Although the impact of large stretches of 4WBM has not been studied in the same detail as that of toehold length, it is nevertheless clear that the kinetics should not have an exponential dependence on the length of the BM domain. Instead, as for a random walk process 4WBM time is expected to be quadratic in the length of the displaced domain, as it is for 3WBM^45^, but with a considerably longer step time^46^. Thus, while 4WBM processes are potentially involved in the displacement reaction and can be crucial for the kinetics of invaders with long toeholds (stable initial binding), it plays less of a role when the accessible toehold is very short.

**Figure 2.**
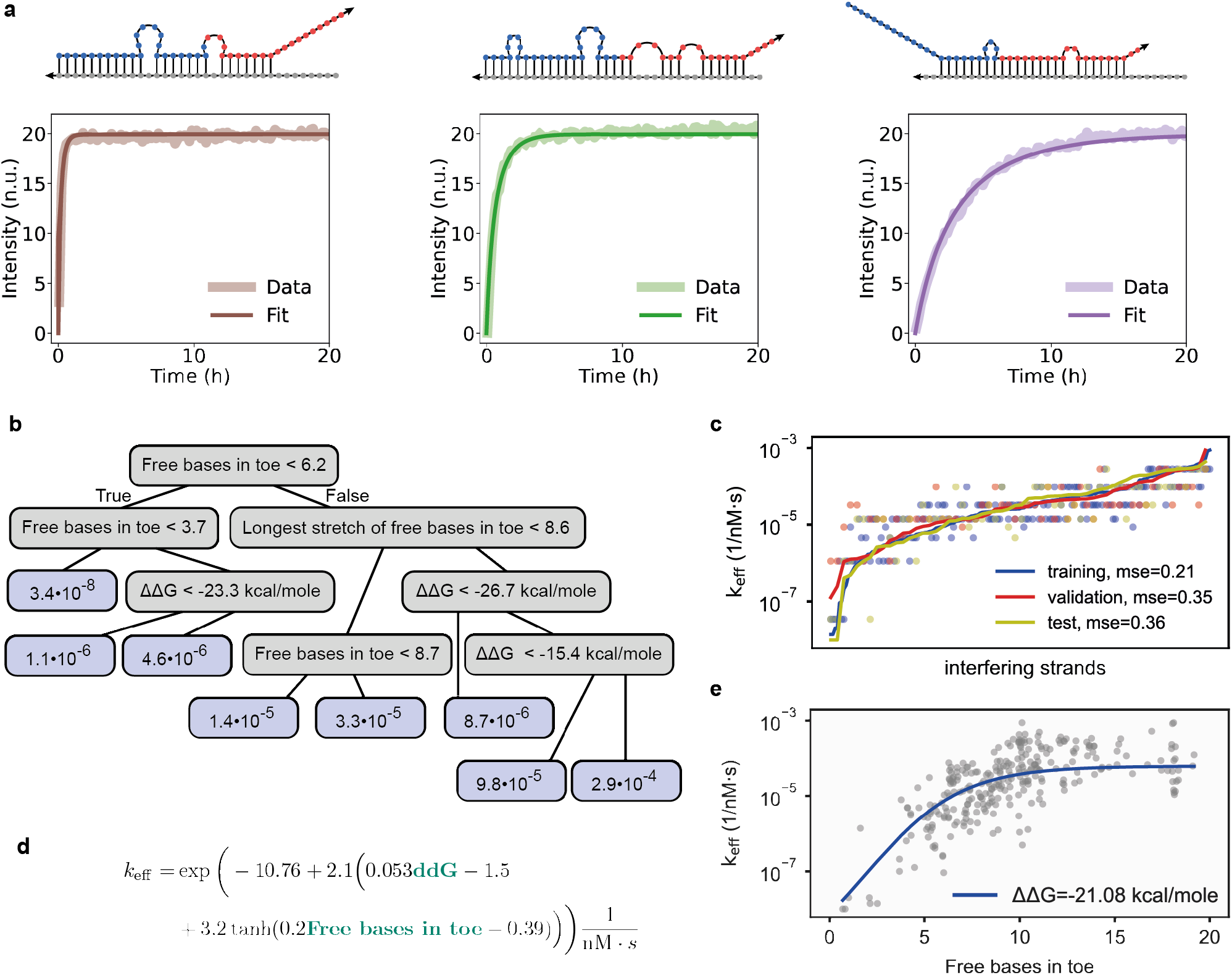
Influence of designed interfering strands on TMSD kinetics. (a) Structure and kinetic curves for three different secondary structures between the invader and an interfering strand as predicted by NUPACK. The colors indicate the toehold (red) and branch migration (BM) domains (blue) of the invader as well as the interfering strand (grey). From left to right, the toehold is more and more occupied but the BM domain less. The kinetics strongly depend on the number of bases bound in the toehold region, concealing the influence of the branch migration domain. (b) Regression tree for the prediction of kinetics based on different structural features extracted from the binding probability matrix between the invader and interfering strands. The blue boxes contain predicted values in units of nM^-1^s^-1^. (c) Prediction of kinetic constants (dots) compared to the measured values (solid line) by the regression tree shown in (b). (d) A simple kinetics prediction model derived by symbolic regression. (e) Predicted value of the kinetic constant (solid line) compared to the measured values (dots) for the model shown in (d) when the ΔΔG value is held constant at the median of the dataset.

To build simple models for the kinetics, we initially explored a relatively large number of potentially useful features from the secondary structure of the invader with the interfering strand. Such features can be derived either from the minimum free energy structure or from the binding probability matrix representing the thermodynamic ensemble (Figure S3). The extracted features include, e.g., the total number of free bases in the invader toehold (Figure S3 and Figure S5a). Figure S4 shows the distribution of some of these features for the two circuits, which are similar overall and cover a range of values that are likely to occur in strongly interacting random pool strands. As expected, the Spearman correlation between kinetics and these features is the highest for those relating to the toehold (e.g., the number of free bases in the toehold), somewhat smaller for those relating to the global secondary structure (e.g., the ΔΔG value), and very small for features relating only to the branch migration (“body”) domain (Figure S5c). In building a simple regression tree, we found that the importance scores for ensemble features are much higher than for MFE features, indicating that they are more useful for kinetics prediction (Figure S5d). We tested a variety of algorithms for their ability to predict kinetic constants based on these features (linear models: Figure S6, regression trees: Figure S7-S10, symbolic regression using Feyn^47^: Figure S11-12, and a neural network: Figure S13-14). A detailed discussion of our findings is given in Supporting Text I.

In brief, we find that a combination of three features, namely the ΔΔG value, the number of free bases in the toehold, and the longest stretch of free bases in the toehold, are sufficient to derive a good estimate of the kinetics for most interfering strands. When using this feature subset, the performance of the more complex neural network is not much better than a small regression tree or a small analytical model, which all are, however, much better than a linear model. For larger feature subsets, symbolic regression and neural networks outperform a simple regression tree, indicating that more accurate predictions require a more detailed consideration of the interfering structure.

Figure 2b shows a regression tree based on three input features and Figure 2c shows an example prediction of kinetics on the dataset. The tree mostly classifies the strands based on the accessible toehold, and then performs finer classification using the ΔΔG values. This is echoed by a simple symbolic regression model, which finds a dependence on toehold length that, for a fixed value of ΔΔG, closely resembles the functional form of the toehold length dependence for a regular strand displacement circuit^45^ (Figure 2d and e, Figure S12e and f). In most cases, the simple regression tree or the analytical models are sufficient to provide a rough estimate of the kinetics, although several strong outliers are observed. For a strongly binding interfering strand, the total number of free bases in the toehold alone already facilitates a good estimation of the kinetics.

### The behavior of random sequence pools

To investigate the influence of large random pools of DNA on TMSD kinetics, we ordered DNA strands with random sequences of defined lengths of 25, 50, 100 and 200 bases (N25, N50, …) and then screened the impact of their concentrations on our measurements. We used two different toehold lengths (10, 20) of our invader. In the absence of background sequences, these two invader strands do not show any significant differences in kinetics. For the following set of experiments, we equilibrated the random pool strands with the invaders for 24 hours before the measurement. After that, we added the reporter strand and started the fluorescence measurement. A sample of the resulting kinetic curves can be seen in **Figure 3**a and b. In contrast to the case of defined interfering strands and invader complexes, fitting of an effective kinetic constant gives less satisfying results, which is perhaps unsurprising as the kinetics are the result of a complex combination of reaction steps. We therefore use the time needed to reach 80% of the final fluorescence value to quantify the kinetics (t0.8). A higher concentration of the random sequence pool leads to slower kinetics and thus more time is needed to reach this level. The concentration dependence of t0.8 results from several effects. First, higher concentrations result in a larger number of transient interactions of unoccupied bases with the random background. Second, the number of well-matching random background strands stably binding more bases of the toehold increases (Figure S17). **Figure 3**a and b also show measurements for different lengths of random pool strands, which present the same total amount N_tot_ of nucleotides (nt) (N_tot_ ∝ strand length × concentration of the random pool). The results suggest a connection between the length of the random strands and their impact on kinetics. For instance, for N25 and an invader with toehold length 10, the difference between 2.5 mM and 5 mM is small, while for the invader with toehold length 20 a higher concentration slows down the reaction considerably. One reason for this behavior could be the higher probability for two different N25 strands interacting with the longer invader simultaneously, as there are more options for binding.

**Figure 3.**
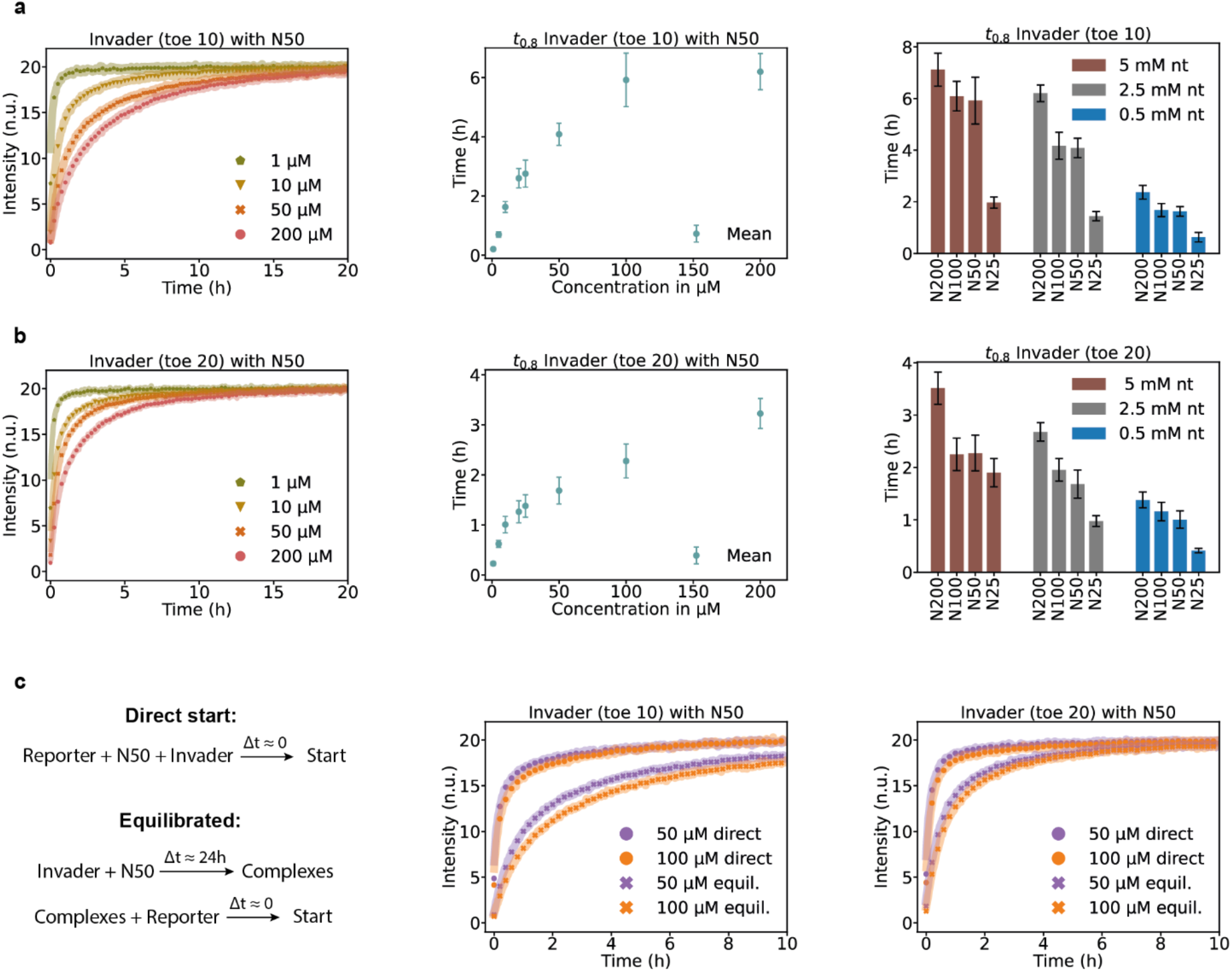
Influence of random DNA pools on TMSD kinetics. (a), (b) Left: Impact of a random pool background on reaction kinetics for invaders with toehold lengths of 10 and 20. The experiments were performed for a random DNA pool strand length of 50 (N50). Middle: Plot of concentration vs. time to reach 80 percent of the final value (t0^.8^) for the two invader lengths. Right: Comparison of t0^.8^ for different lengths of the DNA pool strands but the same total nucleotide concentration calculated as length times concentration. (c) Left: Scheme of the different preparation methods used to measure the kinetic curves. Middle and right: Kinetic curves for the two different preparation methods. The curves for a direct start after mixing the invader with the random pool show fast kinetics in comparison to the measurements after equilibration of invader with the random pool.

Overall, the different behavior of the two toehold lengths in the presence of background sequences can be interpreted in a relatively straightforward way. As our results from individual interfering strands showed, the most important factor influencing the kinetics is the probability of binding enough bases of the toehold to slow down initial binding. For the shorter toehold, only a small number of bases need to be sequestered by interfering strands to significantly reduce the reaction speed as TMSD kinetics become exponentially slower for toeholds shorter than 8 nt. For the longer toehold, it is easier to find partial matches, but a much larger number of bases needs to be bound to significantly reduce the reaction speed. Complete matches are highly unlikely as their probability scales with the length l of the toehold as 4^-l^. The largest effect can therefore be observed for short toeholds in combination with long random pool strands.

In the experiments described so far, the invader had been equilibrated with the random background before the measurement. In typical applications, however, invader strands will encounter the background pool right at the start of the reaction, i.e., without prior equilibration. We therefore also assessed the influence of random strands in experiments, in which all components of the reaction are mixed and the measurement is started directly (**Figure 3**c). Notably, the kinetics for this “direct start” method is much faster, presumably because the invader strands do not have enough time to sample many well-matching strands within the random pool before binding to the substrate.

### Mimicking and modelling the behavior of random pools

In thermodynamic equilibrium (which is approximated by the pre-equilibrated measurements), the concentrations of different complexes can be calculated based on their ΔΔG values. To computationally model the random pool, we sampled 100,000 random strands of defined length (50 nt) and used NUPACK to simulate their interaction with our invader strand. We then split the resulting distribution of ΔΔG values into four energy intervals chosen to cover different regions of interest, of which the first includes strands within the top 1% of ΔΔG values. The last interval includes all strands with less interaction energy than the maximum of the distribution. The boundary between the second and third intervals was chosen to be the median ΔΔG of the first three intervals (Figure 4b). We find that even though the first interval only includes 1% of all strands, in thermodynamic equilibrium almost all invader strands will be bound by such strongly interacting strands (Figure 4c) as long as the random pool concentration is sufficiently high (i.e., when the concentration of these complexes is ∼exp(- ΔΔG/RT)). A feature extraction comparing the whole random pool with the 1% best binders can be found in Figure S18.

**Figure 4.**
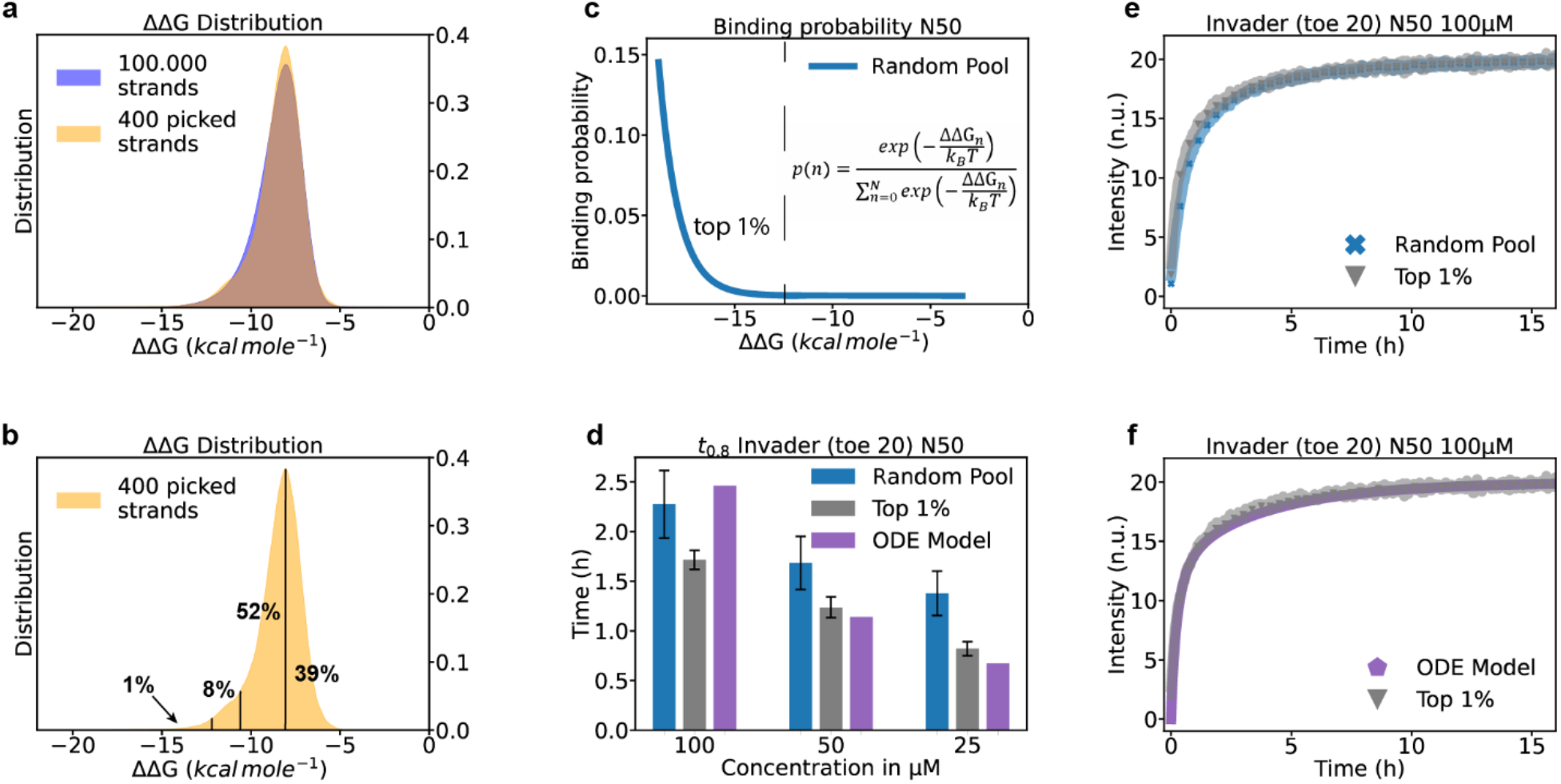
Comparison of a random pool of DNA strands with a pool containing defined sequences (mimic pool). (a) Overlay of NUPACK calculations of ΔΔG values (individual strands to invader-strand complex) results for 100,000 random pool strands with a length of 50 nt and a sampled pool of only 400 strands. (b) The sampled pool is divided into four intervals with different binding strengths, each of which contains 100 strands. The percentages show the contribution of each interval to the total energy distribution. For the following mimic pool measurements, we only used the 100 strands representing the strongest 1% of the random pool. (c) Binding probability versus ΔΔG value of a complex. Strands outside of the upper 1% have almost no probability of binding. (d) t0^.8^ for the random pool, the mimic pool consisting only of the top 1% strands, and the ODE Model. (e) Kinetic curves for the random pool and the mimic pool. (f) Kinetic curves for the mimic pool (top 1%) and the ODE model.

For each of the four ΔΔG intervals, we next sampled 100 different sequences and ordered the corresponding oligonucleotied. By mixing the sampled strands in the correct proportion, we can thus generate reduced pools of strands that recapitulate the ΔΔG distribution of the whole random pool (Figure 4a). We expected that the strongest binding strands would dominate the kinetics and therefore used the DNA pool consisting of only the 100 strands representing the upper 1% of the distribution (referred to as the ‘mimic pool’ in the following) to test whether this pool is already sufficient to recapitulate the reaction kinetics of the full random pool. We adjusted the concentration of the mimic strands in accordance with the random pool concentration we wanted to emulate: for a random pool concentration of 50 µM, the upper 1% of binders would have a concentration of 500 nM, so each of the 100 strands of this pool would have a final concentration of 5 nM in the mixture.

As all of the 100 sequences of the mimic pool are known, we were next interested whether we could find a model that predicts kinetics based on the individual impact of these sequences. Such a model would provide a link between the measurements performed with single interfering strands in Figure 2 to those with the full-fledged random pools in Figure 3. We therefore individually determined the k_eff_ values of the 20 strongest binding strands within the mimic pool. We then used these values to model the kinetics in the presence of multiple interfering strands by calculating initial complex concentrations via the ΔΔG values and treating the reactions for each individual complex as independent apart from the shared reporter complex. A detailed description of the computation is provided in the SI. The temporal change of the concentration of the reporter complex R comprising substrate and incumbent and the individual invader – interfering strand complexes (C_n_) can thus be described as:

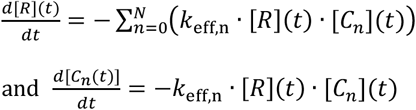

As the initial concentrations are highly sensitive to small changes in the ΔΔG values, we also tested the effect of two different energy models for NUPACK and also allowed for complexes consisting of more than two strands (cf. Figure S19). Comparing the data of the full random pool measurement to that with the mimic pool and to the calculated curve for our selected model supports the hypothesis that only very few of the strongest binding strands of the random pool dominate the equilibrium behavior of the mixture, as the obtained kinetic curves are very similar and the t0.8 is on the same scale (Figure 4d-f). Experiments involving the 100 strands representing the interval with the second best binders show much faster kinetics, confirming these results (Figure S16).

Even though the mimic pool appears to reproduce the overall behavior of the random pool very well, the kinetics of the latter are still a little slower. Furthermore, the mimic pool has slightly slower kinetics than what is predicted by the model at all but the highest concentrations. The comparative slowness of the full random pool may be caused by the invaders not only binding to the best matching strands, but also to the dynamic binding and unbinding of other random pool strands during the search of the invader for the substrate. We have also neglected the influence of binding of random pool strands to the substrate’s toehold.

One should note that the higher the concentration of the random pool and the lower the concentration of the invader strands, the more mimic strands need to be used to obtain reliable results. For instance, in order to mimic the behavior of a 200 µM random pool, 2 µM would represent its best binding fraction and using a mimic pool of only 100 strands then results in a concentration of 20 nM per strand. For experimental invader concentrations of 20 nM or less, the single strongest binder of the mimic pool would therefore bind all of the invader strands and thus completely dominate the reaction kinetics. The fewer strands dominate the kinetics, the more the specific choice of any individual strand will influence the results. In a “real” random pool of length 50 (as opposed to the mimic pool), every strand is statistically likely to be present only once, as the number of possible sequences (4^50^) exceeds the number of actual strands by many orders of magnitude.

### Speeding up displacement kinetics in random sequence pools

For applications, it will be desirable to retain fast and robust strand displacement kinetics despite the presence of a sequence background. One widespread technique to prevent undesired interactions is using a three-letter alphabet lacking one of the bases (A,C,G, or T) for each strand, leading to less stable binding with a sequence background of a different sequence composition (Figure S20). However, this technique also strongly limits the design space and cannot be used for inputs with a fixed (four-letter) sequence. A second technique uses secondary structure within the invader strand itself to prevent strong binding of other sequences, referred to as “internal toehold protection” (ITP) in the following). Similar approaches have already been used by other groups to protect strands from unwanted interactions.^48,49^ Again, this technique limits the design space, as it necessitates potentially unwanted base pairing relationships within the circuit and may be incompatible with external sequence constraints. Ideally, one would like to use an approach that does not constrain the design space, while still making TMSD circuits robust with respect to an arbitrary sequence background.

In order to address this issue, we also pursued a novel approach, which we termed “external toehold protection” (ETP), where an additional strand is bound to the invader with multiple small internal loops. This makes the invader strand less accessible for the nucleic acid background, but still results in a fast displacement rate due to the internal “kinetic toeholds”^16^ provided by the loops within the branch migration domain. Because branch migration in the reverse direction is unlikely once these loops have been bound, they split the random walk process into parts that are completed individually. This greatly speeds up the process as the time scale of the branch migration process is quadratic in the number of displaced bases. Schematics of the internal and external toehold protection techniques are shown in Figure 5a.

**Figure 5.**
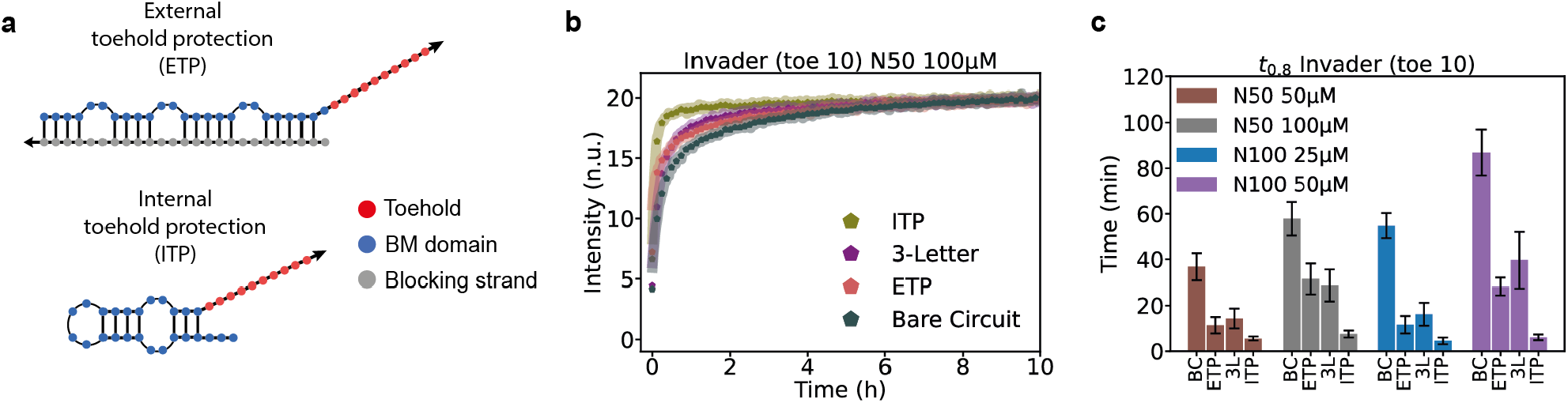
Techniques that speed up TMSD kinetics in random pools. (a) Schematic of two different ways to indirectly protect the toehold of an invader strand by restricting the accessibility of the branch migration domain. (b) Kinetic curves for the different techniques and the bare circuit (BC) in the presence of 50 µM of N100 DNA random pool. (c) Comparison of the t0^.8^ values for the different techniques. All of the tested techniques speed up the reaction kinetics in random pools, although ITP leads to the strongest speed-up by far.

Because the TMSD components are not expected to be in thermodynamic equilibrium with the background nucleic acids in practical applications, we did not pre-equilibrate the invader with the random pool in this set of experiments (cf. Figure 3c), but directly started the experiments after mixing of the constituents. Only in experiments using the ETP technique, the invader was initially incubated with an excess of the blocking protecting strand (Figure 5a in grey) to allow the protective complex to form in advance. A comparison of measurements using the three different techniques with the kinetics of an unprotected invader strand of toehold length 10 is shown in Figure 5b. In these experiments, we chose reporter complexes with the short 10 nt toehold, for which we expected a more pronounced effect on displacement kinetics. All three techniques successfully speed up the reaction kinetics in all random sequence backgrounds tested (Figure 5c), but ITP is found to have the largest impact by far, whereas a three-letter alphabet (in this case the invader strand lacks guanine bases) and the ETP technique show similar, but slower kinetics.

It should be noted that in our case the toehold of the invader for the three-letter alphabet has a higher GC content, which means stronger binding to the random pool strands, but also to the toehold. Nevertheless, the simulations shown in Figure S20 suggest that kinetics should be slightly faster even if the GC content was the same. Several effects might contribute to the differences between ITP and ETP kinetics. First, for the ITP technique less 4WBM is involved, but the toehold is similarly protected from the random pool strands. This is because displacing the first half of the BM domain in the ITP technique automatically removes secondary structure from the second half. A second potential reason for the comparatively large speed-up by the ITP approach is that the incumbent strand necessarily has secondary structure, too, and thus makes its unbinding thermodynamically more favorable.

## Discussion

In this work, we have taken a step towards a better understanding of unwanted cross-reactions with background sequences that can occur in DNA-based chemical reaction networks utilizing TMSD. Such a background could either be present as biological sequences (in a biosensing or an *in vivo* computing context) or as other components of a large circuit. By designing approximately 300 interfering strands with well-defined interactions with the invader strand, we studied the influence on TMSD kinetics caused by binding of external strands to the toehold or the branch migration domain. Even though this in-depth study with feature extractions and measurements of the kinetic curves for interfering strands was performed for only two specific circuits, we believe that the results should be of general use as long as the invader has an approximately uniform distribution of the four different bases.

We derived features from the secondary structure formed between the invader and interfering strands and used the resulting data to build predictors of TMSD kinetics using different machine learning approaches (decision trees, symbolic regression, and neural networks). We find that a small number of parameters, namely the ΔΔG value and two different measures of toehold accessibility - the total number and longest consecutive stretch of free bases – are already sufficient for a rough estimate of kinetics. Although features in the toehold region dominated the overall kinetics, long stretches of bound bases in the branch migration domain also considerably slowed down TMSD. As discussed below, it would be desirable to be able to precisely predict kinetics for any given invader sequence and a list of background sequences. Such a general quantitative model would require a large amount of additional training data over a variety of different circuits.

We next assessed the influence of a random pool of sequences on TMSD kinetics. As expected, longer DNA strands and higher concentrations had a larger impact on the displacement kinetics. **Figure 3**d and e show the different t0.8 for the same total amount of nucleotides but with different strand lengths and concentrations. For N100 and N50, the behavior is almost the same for the same total amount of nucleotides, but the behavior was different for shorter and for longer background sequences. This is likely related to the length of the invader strand itself. For an invader of length 45 nt (such as our toe20 strand) an N25 background sequence cannot possibly bind the whole invader. The different behavior for N200 is harder to explain but might be caused by differences in the synthesis process of longer oligomers, which might also lead to more pronounced sequence biases and thus have an influence on the randomness of the sequences.

We found a large difference in kinetics for two different sample preparation methods. When the invader strand was not given time to equilibrate with the random background sequences, strand displacement was faster by one order of magnitude compared to the equilibrated case. The impact of random interfering sequence is thus highly dependent on the details of the experimental protocol.

In a direct experiment without pre-equilibration, corresponding to most practical applications, a simple one-step circuit will be considerably less sensitive to background sequences than a slower multi-step circuit, in which circuit components which are activated later can equilibrate with the background while the initial stages of the circuit progress. This effect will likely also exacerbate differences between fast and slow circuits in general: a slower circuit, e.g., with a short toehold, will be slowed down disproportionately compared to a fast circuit as it has more time to equilibrate and sample the random background.

In the pre-equilibrated case, our experiments have shown that TMSD kinetics in the presence of a random pool can be faithfully mimicked by a much smaller collection of strands representing only the best matches with the invader within the pool (Figure 4). A straightforward kinetic model assuming a simple superposition of the reactions with the individual strands recapitulates the overall kinetic behavior quite well, supporting the notion that only a small fraction of the background sequences dominates the speed of the reaction. This finding suggests a general method to estimate the kinetics of a TMSD reaction in complex environments: i) find the sequences that are likely to have the largest impact on kinetics, potentially by searching for overall sequence similarity or complementarity with the circuit components; ii) estimate the impact of each individual sequence by a machine learning model similar to the one developed in this work; iii) super-impose the individual estimated kinetics for an estimate of the total impact.

For the non-equilibrated case, where one cannot use structure prediction algorithms to determine, e.g., the distribution of free toehold bases, prediction of the displacement kinetics is considerably more complex, as in this case the circuit components temporarily bind strands of the random pool in a search for the strongest binding partner, while also engaging in the TMSD reaction. Given a model of the equilibration process with the random pool, the reaction speed at each time step could potentially be predicted by a feature-based machine learning algorithm as presented here, yielding an overall predictor of kinetics. Future work should therefore investigate the equilibration process in more detail.

We finally compared three different experimental approaches to speed up SD processes in the presence of a random sequence pool, which ideally should work without any detailed knowledge about the sequence background. As in most real-world applications the circuit components are not expected to fully equilibrate with their sequence environment, we studied the influence of these techniques in direct (non-equilibrated) measurements. All three techniques – one using a three-letter alphabet and two approaches termed internal and external toehold protection (ITP and ETP) – were found to improve kinetics in a random background.

The three-letter alphabet is very simple to use, but too restrictive when using biological sequences as inputs. The performance of the ITP technique – using invader strands with secondary structure for protection - was found to perform best, and thus ITP appears to be the method of choice when the design constraints allow for it. This will likely be the case when there are no external sequence constraints, but ITP might also be applicable when sensing an appropriate subsequence of a longer biological sequence such as an mRNA. As an interesting aside, the success of the ITP approach indicates that when sensing natural sequences using a TMSD circuit, the best input sequences might not be those without *any* secondary structure, as one would naively expect, but instead sequences with an intermediate amount of secondary structure (protecting against interactions with the background) and a highly accessible toehold. The ETP a introduced in this work performs slightly less well than ITP, but can be applied to almost any given input sequence by the addition of a protecting strand with suitable complementarity and is also easier to design than ITP. Thus, both ITP and ETP should be useful approaches to speed up TMSD circuits in complex environments, and the choice of which of the two to use will depend on the specific sequence constraints of the application.

## ASSOCIATED CONTENT

## Supporting Information

The Supporting Information contains Materials and Methods, a Supplementary Text outlining the Machine Learning approaches taken in this work, Supplementary Figures S1-S20, and Supplementary References 1-5.

The data generated in this study and the scripts used to evaluate the data are available for download at Github (https://github.com/loesinghaus/randompooltmsd).

## AUTHOR INFORMATION

## Author Contributions

The manuscript was written through contributions of all authors. / All authors have given approval to the final version of the manuscript. / ‡These authors contributed equally. (match statement to author names with a symbol)

## Notes

The authors declare no competing financial interest.

## ACKNOWLEDGMENT

The authors thank Erik Winfree, Lulu Qian, Chris Thachuk, David Soloveichik and Ho-Lin Chen for insightful discussions as well as all attendees of the DNA28 conference for helpful feedback and criticism of the work.

## For Table of Contents Only

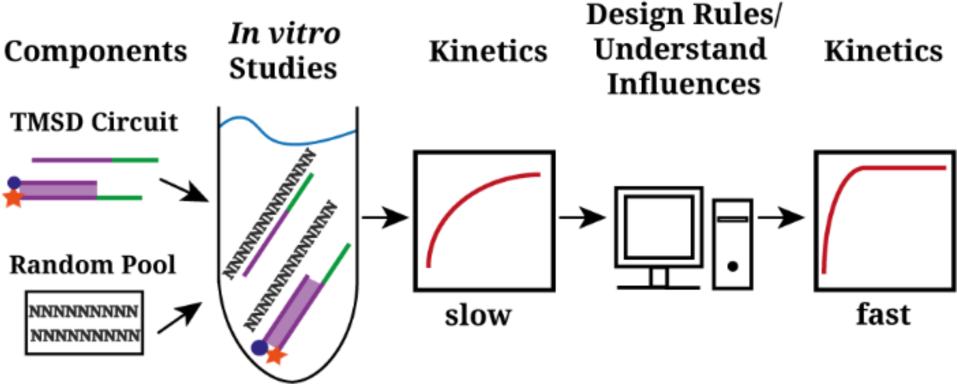

