## Supporting Information for "Toehold-Mediated Strand Displacement in Random Sequence Pools"

#### Materials and Methods

##### DNA design and preparation

We used NUPACK[1] version 4 for all of our designs. The bare circuit design was chosen to be free of secondary structure and to have a mix of all four bases to ensure that our results have a higher general validity. For our reporter system we chose an incumbent labeled with a 5' Iowa Black® RQ and a substrate with a 3' ROX (NHS Ester) fluorophore. All circuit components were ordered at IDT. For the main components (invader, incumbent, substrate) we chose to order them lab ready and PAGE purified. For the set of designed interfering strands (Figure 2) we used a Python program to create randomized MFE structures and NUPACK to design the sequences generating these structures. For these strands we did not choose PAGE purification because less of an influence on the results is expected for possible mismatches and/or synthesis erros, as we used them in a five times excess over the invader. Random sequences were ordered as dried DNA to be able to dilute them to all desired concentrations. For every dilution step and all experiments we used a TE buffer with additional 100 mM NaCl and 12.5 mM $MgCl_2$. For strands with very high concentrations, we added additional $MgCl_2$ to balance the total backbone phosphate charge with magnesium ions. Poly-T strands with a final concentration of 1 µM were added to dilutions below 1 µM to prevent strands from sticking to tubes, tips, and plates. A lsit of the sequences of the used main circuit components can be found under the "DNA Sequences" section.

##### Fluorescence measurements

All fluorescence measurements were performed using a FLUOstar® Omega plate reader by BMG LABTECH and Corning® low volume, non-binding plates. The excitation filter was set to 584 nm and the emission filter to 640 nm. The different experimental constituents were pipetted in the plate, spun down and the measurement was started. Depending on the experiment, a delay between the last pipetting step and the first data point between 30 seconds and three minutes must be considered. For single, very fast experiments (Figure 1c top), we used the built-in syringe of the plate reader to avoid any delay due to pipetting. However, for multiple, parallel experiments or durations of several hours this technique was not be feasible as it takes some time for the plate reader to move the plate from one well to the next, trigger the syringe and make a measurement. During that time no data point for the first well can be taken. Furthermore, the plates cannot be covered and therefore evaporation is much more of a problem. For experiments where we wanted the invader to be in equilibrium with the (designed or random) background sequences, we mixed the invader with the desired sequences and let it incubate for 1 hour in the case of designed sequences and 24 hours for the random background. Non-equilibrium experiments were performed by mixing all constituents and directly starting the measurement.

##### Fitting of kinetic data

The kinetic curves for the bare circuits and the designed interfering strands obtained by fluorescence measurements were fitted using the python packages lmfit and scipy.integrate using the following system of ordinary differential equations:

$$\frac{d[R]_t}{dt} = -k_{\text{eff}} \cdot [R]_t \cdot [C]_t$$

$$\frac{d[C]_t}{dt} = \frac{d[R]_t}{dt}$$

$$\frac{d[I]_t}{dt} = -\frac{d[R]_t}{dt}$$

$[R]_t$: Concentration of reporter complex at time t

$[C]_t$: Concentration of invader-interfering strand complex at time t

$[I]_t$: Concentration of free incumbent at time t
which is directly related to the measured fluorescence intensity

$k_{\text{eff}}$: Effective kinetic constant of the system

#### Mathematical estimates of the sequence space and number of strands

For the different lengths $N$ of our random background strands the number of possible different sequences $Q$ can be calculated as:

$$Q = 4^N$$

For example, with $N = 50$ this leads to

$$Q = 4^{50} \approx 10^{30}$$

different sequences. If we now assume a given concentration $C$ of random strands we can calculate the number of strands $S$ in our random pool using Avogadro's number $\mathrm{N_A} = 6.022 \times 10^{23} \frac{1}{\mathrm{mol}}$ and our volume $V$ per experiment:

$$S = C \times V \times \mathrm{N_A}.$$

In our experimental setup we have typical values of $C = 100$ µM and $V = 20$ µl which leads to

$$S = 100 \times 10^{-6} \frac{\mathrm{mol}}{\mathrm{l}} \times 20 \times 10^{-6} \mathrm{l} \times 6.022 \times 10^{23} \frac{1}{\mathrm{mol}} \approx 1.2 \times 10^{15}$$

strands in the solution, which is much less than Q, indicating extreme undersampling of the sequence space for $N = 50$. For the shorter $N = 25$ it is theoretically possible, however, that every single sequence actually is present in the experimental random pool as we have

$$Q = 4^{25} \approx 1.1 \times 10^{15}$$

different sequences but the same number of strands $S$.

#### Computation of equilibrium concentrations of multiple complexes containing the invader strand

For the models in Figure 4 and Figure S19, we had to determine the equilibrium concentration of complexes in the presence of multiple interfering strands binding a single invader. The ΔΔG value for every invader-interfering strand complex was obtained via NUPACK and used to calculate the concentration of each complex by solving a system of equations with a python integrated least squares method under the assumption of a complex size of at most 2. Calculation of the concentrations proceeded by solving a system of N coupled non-linear equations of the following type:

$$\frac{[AB_n]\rho_{\mathrm{H_2O}}}{([A]_0 - \sum_{n=0}^{N}[AB_n])([B_n]_0 - [AB_n])} = \exp\left(-\frac{\Delta\Delta \mathrm{G}}{k_B T}\right)$$

with

$A$ = invader strand

$B_n$ = interfering strand n

$AB_n$ = complex containing A and $\mathrm{B_n}$

$\Delta\Delta\mathrm{G} = \Delta G_{AB_n} - \Delta G_A - \Delta G_{B_n}$

$\rho_{\mathrm{H_2O}} \approx 55.3 \frac{\mathrm{mol}}{\mathrm{l}}$ at 29°C

Here, every interaction of some interfering strand $B_n$ with the invader $A$ leads to a complex reducing the concentration of the invader $A$, which couples the equations for the N different interfering strands.

#### Computation of displacement kinetics with multiple complexes

We numerically solved a system of ordinary differential equations based on the initial concentrations given by the ΔΔG values and the assumption that the total reaction is simply given by a sum of individual second order reactions with a single $k_{eff}$ value each that run concurrently and are independent apart from competition for the same substrate. Under these assumptions, the concentration change of the reporter complex R comprising substrate and incumbent and the different invader – interfering strand complexes ($AB_n$) can be described by

$$\frac{d[R](t)}{dt} = -\sum_{n=0}^{N}\left(k_{\mathrm{eff,n}} \cdot [R](t) \cdot [\mathrm{AB_n}](t)\right)$$

and $\frac{d[\mathrm{AB_n}(t)]}{dt} = -k_{\mathrm{eff,n}} \cdot [R](t) \cdot [\mathrm{AB_n}](t)$.

#### The distribution of random pool ΔΔG values and probability distribution of complex formation

We used NUPACK simulations to obtain ΔΔG values for the interaction of different random strands with our invader. To generate a statistically valid energy distribution, we simulated the interaction of 100,000 different random strands with the invader (Figure 4a). To determine which of these random strands are most relevant to the kinetics, also the probability p(n) is required of the invader being bound to the strand n when the entire pool is present. We approximated this probability by assuming a Boltzmann distribution

$$p(n) = \frac{\exp\left(-\frac{\Delta\Delta \mathrm{G}_n}{k_B T}\right)}{\sum_{n=0}^{N}\exp\left(-\frac{\Delta\Delta \mathrm{G}_n}{k_B T}\right)},$$

which is valid when the concentration of invader strands is much lower than the concentration of random strands.

#### Normalization of fluorescence measurement data

The fluorescence data was provided by the software in arbitrary units (a.u.). We used two different methods to normalize the data. For both methods, the background signal of a well containing only buffer was measured and subtracted from each measurement. For the interaction with the designed strands (Figure 2), we measured the maximum fluorescence intensity for the basic strand displacement circuit without any interfering strands and, after subtraction of the background, this value was set to 20 because of the 20 nM invader we used. All other measurements were normalized to this measured fluorescence value, because not all of the experiments including interfering strands reached the final value as they were extremely slow. The second method, which we used for interaction with the random pools and mimic pools, was to take the last 10 values of the measurement data in a.u., subtract the background, and set the mean fluorescence value to be 20. This was feasible because we expected the displacement process to run to completion after the 24h measurement time as there are no complexes with all toehold bases occluded (cf. Figure S17c).

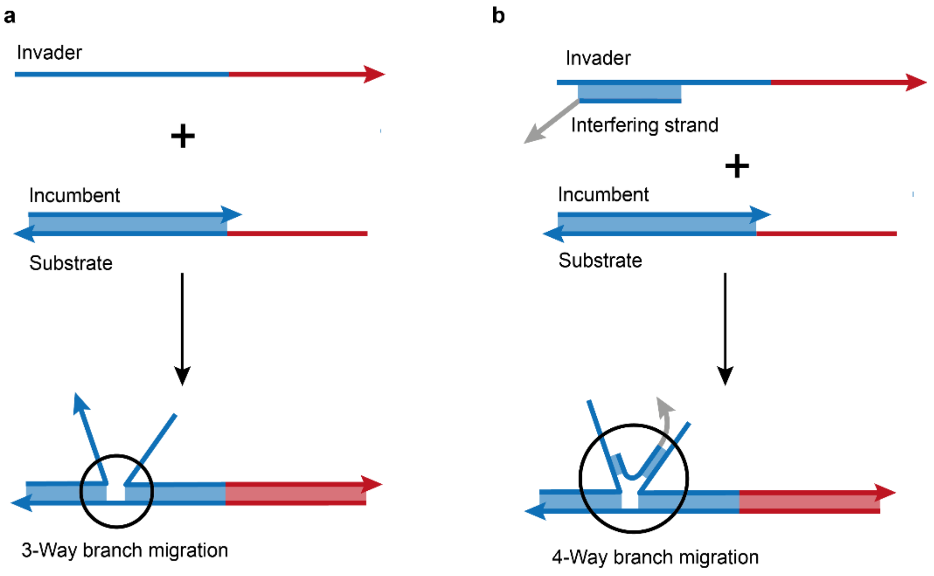

**Figure S1. 3-way and 4-way branch migration. a)** The invader is single-stranded with a red toehold and a blue branch migration domain. After the initial binding to the toehold of the substrate strand, the branch migration process takes place and the incumbent and invader compete for binding in a 3-way process. **b)** A part of the invader's branch migration domain is bound to an interfering strand. This subsequently leads to a more complex and, in general, slower 4-way branch migration process as not only the substrate strand but also the interfering strand is partially complementary to the incumbent and invader strand.

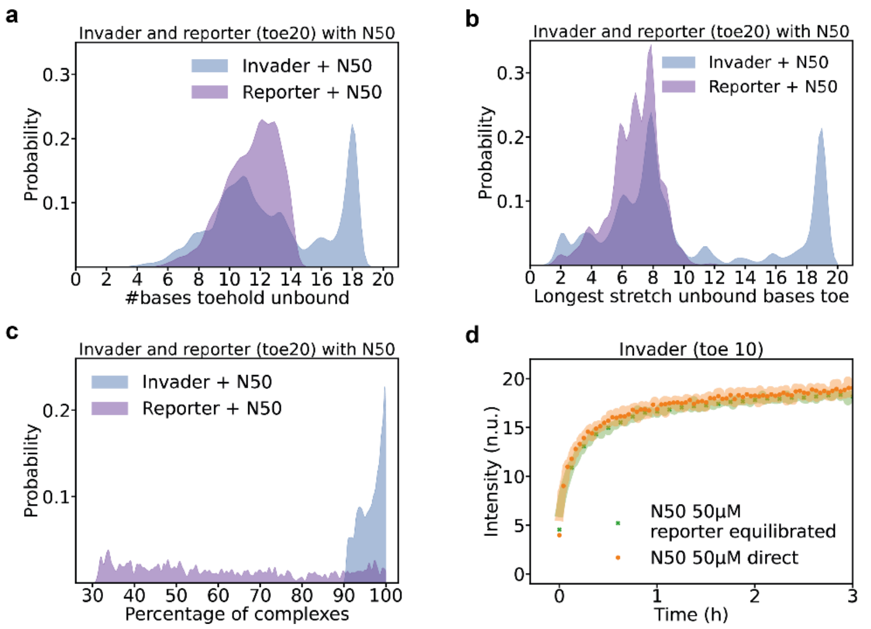

**Figure S2. Interaction of the reporter toehold with a random pool.** Interaction between the N50 pool with the invader and reporter toehold. For subfigures a)-c), only the top 1% of binding strands of the random pool were taken into account for feature extraction, as they dominate the displacement kinetics. For better visualization of the histograms, kernel density estimation using an Epanechnikov kernel was chosen. **a)**, **b)** The number of unbound bases in the toehold has a wider distribution for the invader with a higher percentage of complexes having almost no bases bound in the toehold, which is likely an artifact of considering only complexes of size 2, but the number of complexes having only a few unbound bases left in the toehold is larger, too. The same is true for the longest stretch of unbound bases in the toehold. **c)** Probability of complex formation depicting the ratio of complex to its individual components in thermodynamic equilibrium. For the interaction of the invader and a random strand of the top 1%, complex formation is very probable and the toehold will most of the time be occluded. The interaction of the reporter and a strand of the pool complex formation is less probable, however, which means that often the toehold will not be occluded in the way that is predicted by the base pairing probability matrix, as it only shows the probability in case the complex has actually formed. **d)** Orange: Kinetics measurement of invader with toehold length 10 and a random N50 pool with 50 µM concentration. The constituents were pipetted together and the measurement started immediately. Green: Same setup but this time the random pool was equilibrated for 24 hours with the reporter.

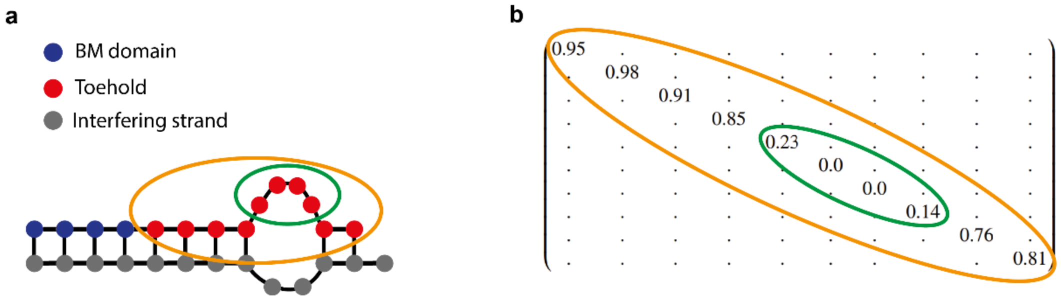

c

#bases toehold unbound = #bases toehold - ∑ binding probability

#bases toehold unbound = 10 - (0.95+0.98+...+0.76+0.81) = 4.37

Longest stretch unbound bases toe = #free bases MFE - ∑ binding probability

Longest stretch unbound bases toe = 4 - (0.23+0.0+0.0+0.14) = 3.63

**Figure S3. Different possibilities for feature extraction.** One can user either the **a)** minimum free energy (MFE) structure or **b)** the probability matrix. The matrix depicted in b) is just an excerpt of the complete matrix and the diagonal entries display the probability of one of the bases of the strand of interest depicted in a) being bound, starting at the 5' end. The probability matrix gives a more complete picture of the secondary structure than the MFE structure, where one can only count bound and unbound bases. For further analysis, we mostly used the ΔΔG of the complex which can just be calculated as well as the number of bases unbound in the toehold region and the longest stretch of unbound bases in the toehold region. **c)** General calculation of the two features as well as corresponding example calculations for the complex shown in a).

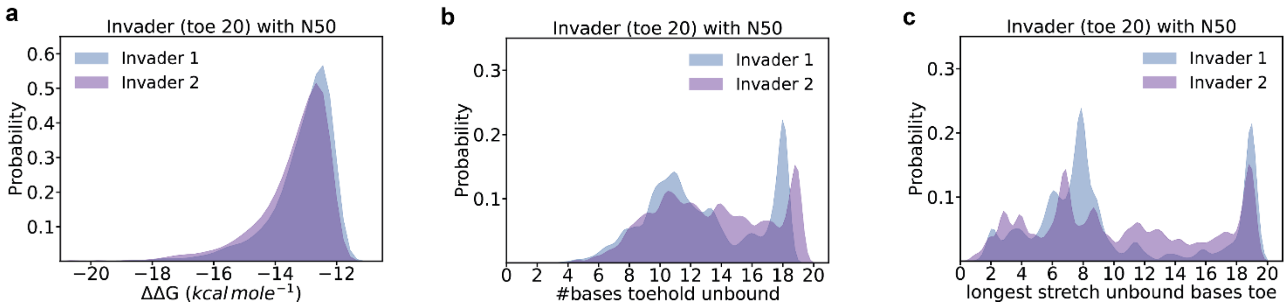

**Figure S4. Feature extraction for two different invader strands of the same length but with different sequences.** For the two invader strands for complexes with designed interfering strands we extracted **a)** the ΔΔG distribution, **b)** the number of unbound bases in the toehold domain, and **c)** the longest consecutive stretch of unbound bases in the toehold domain. We chose to cover a range of interaction energies where the interfering strands bind stably to the invader as such strands will be most relevant to the kinetics.

### Supplementary Text I – Machine learning-based analysis of interfering strands

#### Introduction

Here, we examine data from interfering strands for two TMSD circuits (Figure S4) to understand how different interactions between interfering strands and the invader alter kinetics. As the amount of data is rather limited, we do not train an end-to-end model based on the sequence or even on the probability matrix as output by NUPACK. Instead, we heuristically extract relevant structural features from the latter and test different algorithms for their ability to predict kinetics based on these features. Apart from reducing the required amount of data, the features are also readily interpretable and thus serve to gain intuition about the impact of different kinds of interfering strands. The resulting models do not predict kinetics for arbitrary interfering strand sequences as certain types of structures are not well-represented in the training data. For example, the training data do not contain any strands that do not bind to the invader strand at all due to a large ΔΔG (ddG) value, and would thus not be predicted correctly. As we will see, other types of interfering strands, such as those derived from the top 1% of random binders, also sometimes contain structural features that are not represented in the training data. The data nevertheless gives insight into the general mechanisms by which interfering strands slow kinetics. We further suspect that the same features and algorithms could be used to predict kinetics for arbitrary interfering strands by increasing the size and diversity of the training data.

#### Selecting salient features

We derived 28 features derived from the secondary structure of the interfering strands in complex with the invader (Figure S3, S5A). We tried to construct the features in a way that minimizes dependence on the precise structure and sequence of the invader strand. Thus, we generally considered free bases in the toehold and bound bases in the body domain. Most features can be calculated in two different ways: from the MFE structure or as an ensemble value from the base-pairing probability matrix (Figure S3). Certain other features one could imagine using, such as the number of free AT bases in the toehold, are trivially computable from the number of free bases in the toehold and the number of free GC bases in the toehold and are therefore here not considered separately.

As shown in Figure 2 and S4, we measured the impact of interfering strands for two different circuits. In the following, data from circuit one and circuit two are pooled to function as a single data set unless otherwise mentioned, giving us a total of 290 data points (198 data points from circuit one and 92 data points from circuit two). Figure S5B shows the Pearson correlation between the different features. The MFE features and the ensemble features, insofar as both are computed, are highly correlated. There are three types of features: global, toehold, and "body". Within each type of feature, the correlation is highly feature-dependent, indicating that a combination of several features from each type might be required. There is limited correlation between features of different types. Figure S5C shows the Spearman correlation between the features and the logarithm of the kinetic constant. In the following, all analyses will be performed on the logarithm of the kinetic constant. As expected, the Spearman correlation between toehold-type features and the kinetic constant is much larger than for body-type features. The correlation for global-type features is less than for the best toehold-type features (i.e., the free bases in the toehold and the longest stretch of free bases in the toehold), but still rather high overall. Across the board, ensemble-type features show a slightly but consistently larger correlation than MFE-type features, indicating that they are probably more useful for kinetics prediction.

Next, we wanted to determine which of our features are most useful for understanding the kinetics. We used scikit-learn (version 1.1.2) to build a regression tree classifier for our data based on all 28 features without pruning of the tree and extracted the importance scores, i.e., the mean decreases in impurity (Figure S5D). Because we have a relatively small number of data points (especially at very small kinetic constants), the precise division into the train, validation, and test sets has a strong impact on the results, which is why we use ten different random seeds for assigning data points to the splits. Here, 60% of the data was assigned to the training set and 40% to the validation set for consistency with later work. First, we observe that the ensemble values are universally preferred to the MFE values, in line with the higher correlation observed in Figure S5C. The number of binding blocks is essentially never used. We therefore discard the binding block features and all MFE features in the following. Second, the number of free bases in the toehold is the most important feature, which is then followed by a global feature such as total bases bound or the ddG value. This seems reasonable looking at the correlations: The number of free bases in the toehold has the highest Spearman correlation. The ddG value/number of total bases bound also show a high correlation with the kinetic constant and, unlike other toehold features, only very limited correlation with the number of free bases in the toehold. In the following, we will try to see which of the remaining features are necessary to understand the kinetics. Based on the data presented thus far, we expected to require at least two features: One toehold-type feature, most notably the number of free bases, and one global-type feature such as the ddG value or the total number of bases bound.

Repeating the importance score analysis using only the ensemble features (Figure S6A) leads to similar results as for the previous analysis. We normalized the features to mean 0 and variance 1 and performed a principal component analysis (PCA) on our data (Figure S6B). The top three components explain most of the variance in our data, but are probably not useful for fitting it as they have a relatively low Spearman correlation with the kinetic constant (Figure S6C).

As a first approximation, we tried fitting a linear regression model to the kinetic constant (Figure S6D), which we shifted to zero mean and unit variance for all fitting tasks for comparability (i.e., the mean squared error (mse) is always calculated on logarithmic kinetics data with zero mean and unit variance). The model performs poorly ($R^2$=0.46) at fitting the data. From now on, we will generally use the mse to assess the performance of models. For a linear model, using only two variables (ddG, free bases in toe) achieves the same performance on the validation set as using all ensemble values (Figure S6E). A test set was not necessary in this case as the model was not fine-tuned on the validation set. The poor performance of the linear model is unsurprising as it would not even be able to fit the expected dependence of kinetics on toehold length for a regular strand displacement circuit[2].

**Regression trees for kinetics prediction**

To create easily interpretable kinetics models, we produced regression trees for different choices of input features. We split the data into 60% training data, 20% validation data, and 20% test data. The trees were pruned using minimal cost-complexity pruning to create a series of trees with a steadily decreasing leaf number. Of these trees, the one with the lowest error on the validation set was chosen. The performance was evaluated on the validation set, the test set, and out-of-distribution data, wherein the latter consists of individual measurements of twenty strands from the 1% mimic set (see main text).

As the importance score measure based on mean decreases in impurity can be biased[3], we tried to identify the most relevant features by dropping each feature individually and assessing the impact on the performance of the resulting regression tree on the validation and test sets. Across the board, dropping a single feature did not lead to any obvious increase in prediction error. This may initially seem surprising given the high importance score calculated for the number of free bases in the toehold (Figure S5D). Because some toehold-type features are highly correlated (Figure S5b), this feature can, however, be replaced by a combination of other toehold-type features when dropped. Therefore, we instead made educated guesses about potentially useful subsets of the ensemble features and tested their performance for prediction (Figure S7A). The error shown is the standard error of the mean (based on ten different assignments of data to train/validation/test sets).

For this data, as for the linear model, there is virtually no benefit in using, e.g., all values over just the ensemble values (model A versus B), or, as far as the validation and test sets are concerned even just two parameters (model B versus model H). In the out-of-distribution data, there does seem to be a benefit in using three parameters rather than just two (model G versus model H). Thus, the first conclusion is that, for a regression tree, the use of more than three of our features does not improve prediction for the amount of data we have available. Using features of different type (i.e., ddG and free bases in the toehold [model H]) is superior to using two features of the same type (free bases in the toehold and longest stretch of free bases in the toehold [model I]). The ddG value and the total number of bases seem to work approximately equally well (model H vs model K). The total free bases in the toehold works approximately as well as the longest stretch of free bases in the toehold (model H versus model J). There is some improved performance for the out-of-distribution set for the latter, which, however, doesn't really reflect "insight" by the model (see below). Use of only a single parameter performs far worse than using two parameters (models L and M).

Thus, for the regression tree, a combination of at least two parameters, one of global-type and one of toehold-type, seems to be the minimum for a decent prediction. As will be explored later, this is not because two such parameters fully describe the problem, but mostly because more data would be necessary to reliably fit regression tree models for a much larger number of features. The body-type features are not used explicitly here – it should be noted, however, that the total number of bound bases in the body domain can be easily calculated from the free bases in the toehold and the total number of bound bases. They can therefore be considered indirectly even when their corresponding features are not used. Figure S7B shows predictions for a specific choice of data split. The tree for model G appears to represent the data relatively well.

Figure S8 shows some of the fitted regression trees. To understand how kinetics are estimated by the trees, we will traverse the tree for model H as an example. The tree first sorts the data into slow and fast strands according to the toehold length (toe < 6.2). Then, it checks if the invader has essentially no accessible toehold at all (toe < 3.7). In this case a very slow reaction can be expected. If the toehold is somewhere in between, very strongly bound strands (ddG < -23) are somewhat slower than more weakly bound strands. This could represent underlying structural features such as AT vs GC binding or many consecutively bound bases. On the faster branch, the tree again starts by dividing according to a very accessible toehold (toehold > 9.5) or an intermediate accessibility. The former likely corresponds to the length above which activation kinetics saturate for most strands. This interpretation is underlined by a different marginal model based on symbolic regression (see below). Then, there are again a number of finer divisions according to the ddG value, which likely represent binding structures in the toehold and bases bound in the body domain.

Looking at out-of-distribution data (Figure S9A), we find that the decision tree for model G usually performs well, but has several severe outliers. Model H, which uses fewer parameters, performs poorly on this data, for reasons that will be explained shortly. Model J, which uses the longest stretch of free bases in the toehold, performs somewhat better, though mostly because it happens to guess values closer to the average in specific areas. To see why model G sometimes fails, we examined three outlying data points (labelled 1, 2, and 3) individually.

The first data point (1) is one that could, in principle, be correctly predicted by the tree given our chosen parameters, but it is not because there are no similar data points in the training data. Specifically, it has a very large number of free bases in the toehold (9.84), but a small longest free stretch of bases. Due to this large "free" toehold, it is automatically classified as relatively fast (i.e., it traverses the tree on the very right), as no strand with such a high value in the training data has slow kinetics. This example shows why simply considering the total number of free bases in the toehold (model H) cannot be sufficient and that the largest stretch of free bases also needs to be considered (model G). In fact, the calculated longest free stretch is still longer than one would think based on the shown structure. The area considered for the longest stretch of free bases is still based on the MFE, and the resulting number can thus depend critically on whether or not an individual base is open or closed in the MFE. Here, slight differences in the energy model between the online and offline versions of NUPACK lead to a closed base towards the end of the toehold for the latter. This excessive sensitivity to the energy model is undesirable and suggests that, when fitting a model with more data, it might be better to calculate the longest stretch of free bases somewhat differently, e.g., by a probability-weighted average over the 95$^{th}$ percentile of probable structures. Ideally, this should be combined with an average over different energy models or even different algorithms. We do not implement this change here because we would anyway require more training data for it to be effective.

While the first data point might be correctly predicted by model G with more training data and adjusted calculations, the second data point (2) can never be correctly estimated by it. This is because the underlying structure has a large number of free bases in the toehold, a large longest stretch of free bases in the toehold, and a relatively large ddG value. The kinetics are nevertheless slow because a large number of bases are bound in total. Thus, a model that predicts this data point should probably also include the total number of bound bases. Note that, given the constraints of our training data, including this feature is insufficient due to a lack of similar training examples.

The third data point (3) is predicted to be much slower than it actually is. The slow prediction stems from its very small free toehold, both in terms of total free bases and in terms of the longest stretch of free bases. There is, however, a large stretch of free bases directly adjacent to the toehold, which might nevertheless allow the invader to bind the reporter there. Such behavior could be captured by a parameter similar to the "bound bases in transition area body" feature.

Thus, the failures of prediction could be ameliorated by slightly changing the way some of the features are calculated and then including such extra features in the prediction, leading to a total number of perhaps five features in a potential improved model (e.g., ddG, total number of bases bound, total free bases in toe, longest stretch of free bases in toe, free bases in transition area body). However, properly fitting a tree to these features would require additional training data and is therefore left to follow-up work.

We next tested what happens when we only use data of circuit one (70% training data, 30% validation data) for fitting and use the data of circuit two as test data (Figure S10). The results are largely the same as for pooled data in terms of conclusions for chosen features. In general, models G and H seem to generalize well from circuit one to circuit two, having almost the same error on the validation and test set. This indicates that the precise sequence of the invader strand is not highly relevant to kinetics prediction using our engineered features, indicating that measuring a small number of circuits is likely sufficient to draw general conclusions about the impact of interfering strands, and that our problems with prediction of the out-of-distribution data indeed stems mostly from a lack of similar structures in the training data.

**Interpretation using symbolic regression**

We also tried using symbolic regression using Feyn[4] (version 3.0.2). Feyn generates analytical model functions for output variables based on training data. The idea here is thus to generate simple models for kinetics with a similar number of parameters as for the regression tree, thus retaining easy interpretability, while potentially improving prediction performance. The train/validation/test split was the same as for the regression trees. We used the auto run feature with a mean squared error loss and the Bayesian information criterion (BIC) for model selection. Figure S11A shows the performance for models B, G, and H. The performance is indeed superior to that of the regression trees. Due to restrictions on complexity enforced by the BIC, only a relatively small number of features is actually used for model B even when all ensemble features could be used as an input, in principle. Figure S11B shows an example for model B, which we randomly chose for illustration, but nevertheless has some interesting features. One such feature is that it adds the bound bases in the transition area toe to the free bases in the toehold, although with a much smaller factor (0.52 vs 0.12). This is precisely what we diagnosed to be the cause for model G outlier 2 in Figure S9A: Many free bases adjacent to the toehold domain could potentially support an otherwise short toehold. Figures S11 C and D show concrete predictions. Generally, the outliers in the out-of-distribution data remain outliers.

A much simpler model using only two parameters (Figure S12A) is useful for gaining an intuition for the kinetics. The number of free bases in the toehold is passed through a nonlinearity, and then simply added to the ddG value. When holding ddG constant at the median value in the data set, a plot over toehold lengths (Figure S12B) yields a plot reminiscent of the expected behavior of regular strand displacement systems[2], although we plot the ensemble value of free bases in the toehold rather than a coherent stretch of free toehold bases. This indicates that, on the average, the number of free bases in the toehold plays a similar role as the regular toehold length does for "clean" strand displacement systems. Figures S12C and D shows specific predictions. Unlike for the fully linear model, this very simple mathematical model correctly predicts the behavior of very slow strands. A similar interpretation can be applied to the longest stretch of free bases in the toehold (Figure S12E). Here, there is an inflection point around 6 free bases, which is where kinetics starts to saturate for regular toeholds in in vitro strand displacement systems. Unlike for these systems, the kinetics continue to slowly become faster afterwards.

##### A neural network for kinetics prediction

To estimate how much information is contained in our training data, i.e., to find how much is missed by the regression tree, we tried fitting a simple fully-connected neural network to it (Figure S13A). We used Pytorch (version 1.13.0) with Pytorch Lightning (version 1.6.4) to train our models. The network consists of a series of fully-connected layers with 30 nodes with ReLU activations. Models were trained for 250 epochs. Making these layers smaller for smaller input sizes did not decrease the validation error, which is why we kept the size constant over the different feature subsets. We used a dropout value of p=0.25 for all layers based on a parameter screen. The network has a total depth of eight layers. Because we noticed a decrease in performance with increasing layer number for a regular multilayer perceptron, we introduced skip connections for the layers in the middle of the network[5].

Figure S13B shows the performance across the different models. The performance for very simple models (G, H), is largely the same as for the regression tree, indicating that not a lot of information is missed by the regression tree models. For a larger number of inputs (see especially model B, which contains all ensemble features), the performance is markedly improved. While the difference might not seem like much, one should keep in mind that the error is calculated on a logarithmic scale. Figure S13C shows predictions for particular networks. The distribution of points for model B is close to the true values with relatively few significant outliers. Thus, the neural network can extract more information from the large number of inputs than the regression tree. This translates to a superior performance on the out-of-distribution data (Figure S14).

##### A comparison between prediction methods

Figure S15 compares the performance of regression trees, Feyn models, and the neural network for the most important feature subsets. This is not meant as a direct comparison of these fundamental approaches, as the former two, by using a single regression tree rather than, e.g., XGBoost for the tree-based algorithm and the BIC for Feyn models, were chosen explicitly for a small model size and easy interpretability rather than minimal prediction error. Across the board, Feyn and the neural network perform better than the regression tree, although the difference is small for the models with few parameters (G, H). For these models, the performance of Feyn and neural networks is virtually identical, although the former uses far fewer parameters. For the larger feature subset (model B), the neural network performs better than symbolic regression, which performs better than regression trees.

In conclusion, we find that a relatively small number of parameters (ddG, total free bases in the toehold, longest stretch of free bases in the toehold) are sufficient to allow for a relatively good estimate of kinetics even using a relatively small regression tree or analytical model. This is fortunate, as these values are very easy to calculate and can essentially be estimated by simply looking at the base-pairing probabilities of a NUPACK structure prediction. Thus, the regression trees and analytical models can serve to build intuitions about kinetics. To get a significantly more reliable prediction of kinetics requires more features (e.g., total bases bound, free bases in the transition area), more training data, and might best be achieved by a neural network-type approach. Ultimately, a very precise prediction would probably have to be based on an end-to-end model using either the base-pairing probability matrix or even just the primary strand sequences.

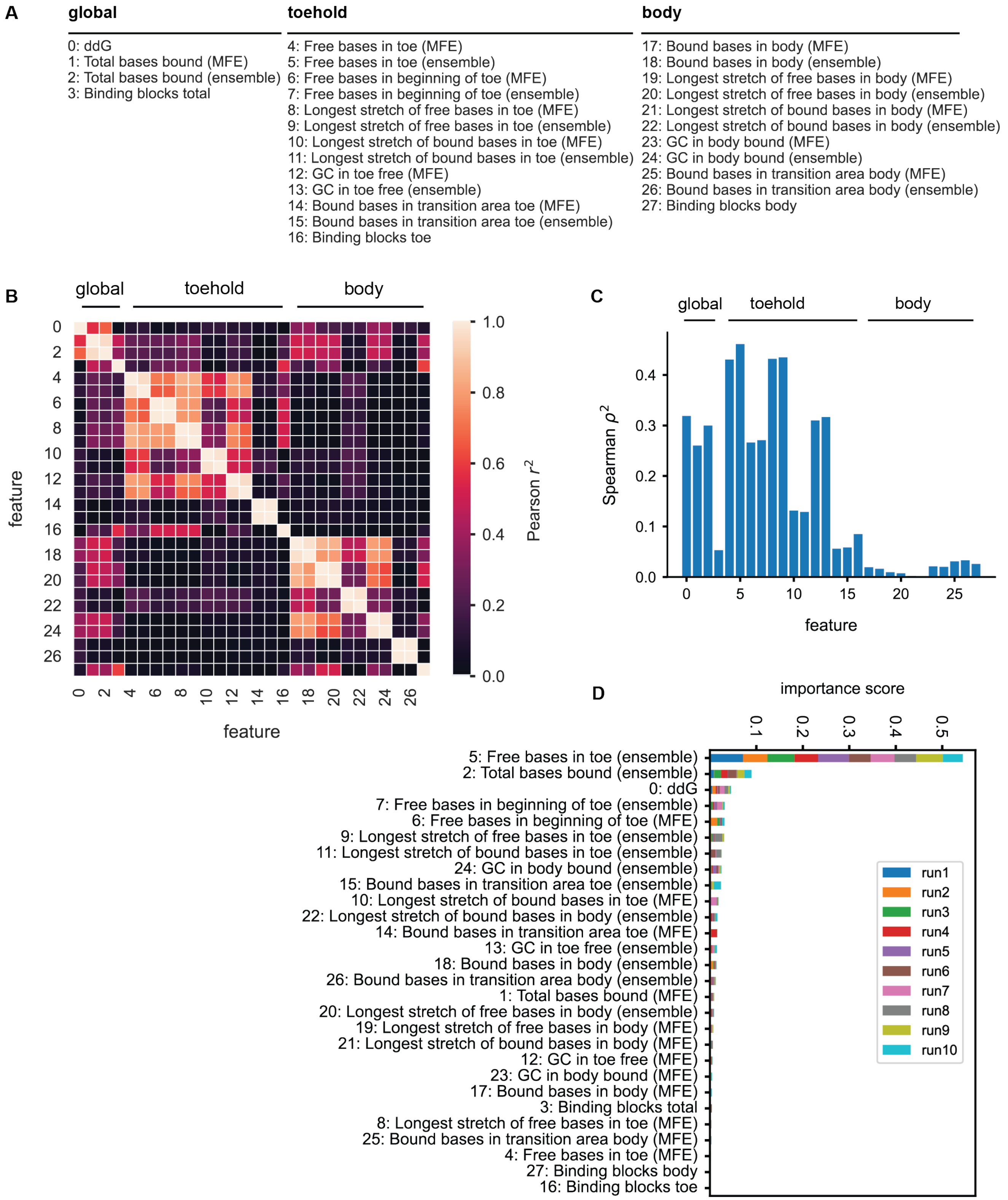

**Figure S5. Choice of interfering strand features. A)** All features that are considered in the evaluation of the secondary structure of the interfering strands and the invader strand for kinetics. **B)** Square of the Pearson correlation between the different features. **C)** Square of the Spearman correlation between each individual feature and the logarithm of the kinetic constant. **D)** Importance score for each individual feature for a non-pruned regression tree based on the mean decrease in impurity for 10 different train/validation splits.

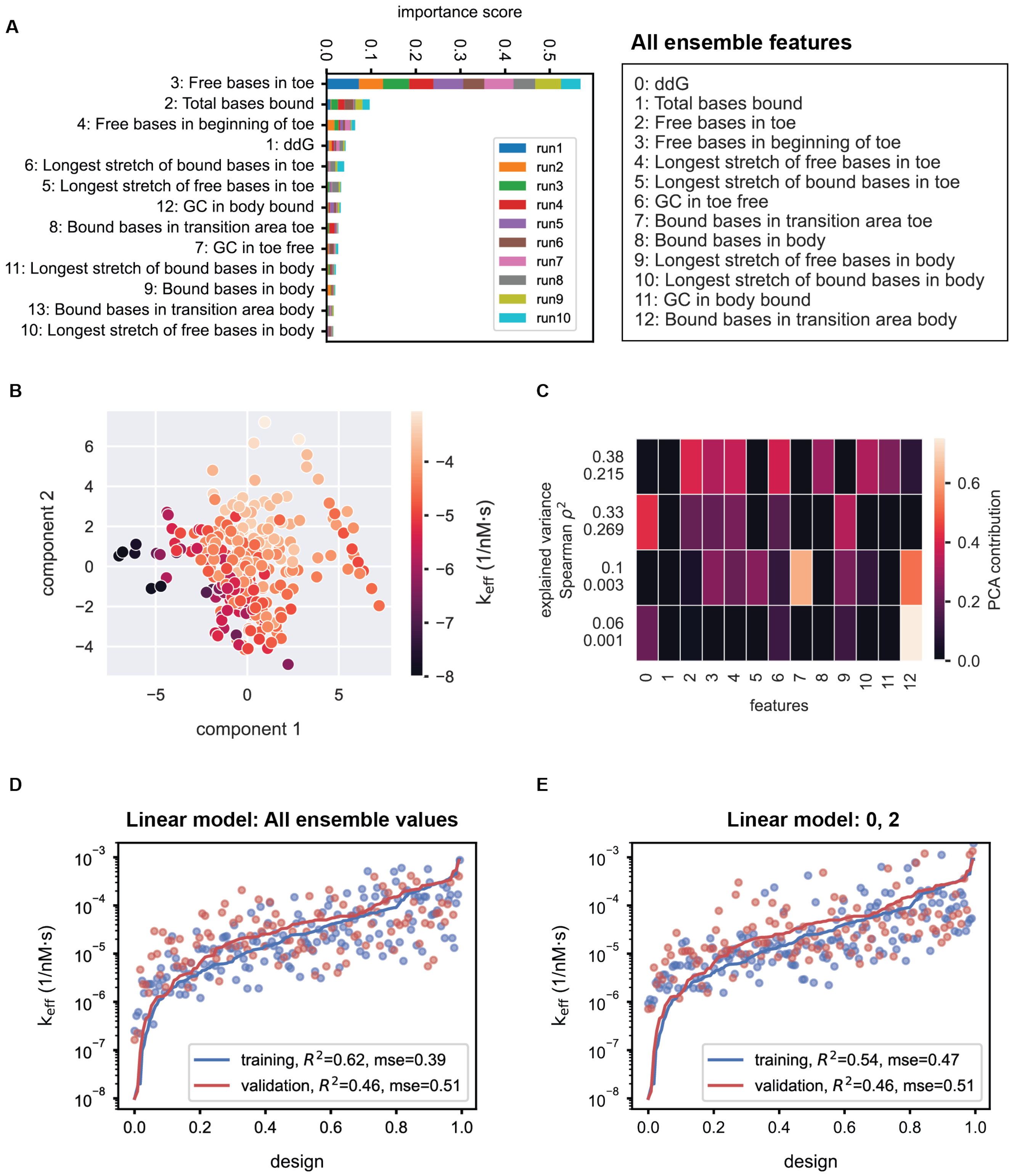

**Figure S6. Linear analysis of interfering strand features. A)** Importance score for each individual feature for a non-pruned regression tree of ensemble features based on the mean decrease in impurity for 10 different train/validation splits. **B)** PCA of the ensemble values for all strands in the two measured circuits. **C)** Contribution of each ensemble feature to the PCA axes. The explained variance of each axis and its Spearman correlation with kinetics is shown on the y axis. **D, E)** Linear regression model based on all ensemble values (D), or only on two ensemble values (E) for the prediction of kinetics. solid line=true value, dots=predicted value, mse=mean squared error

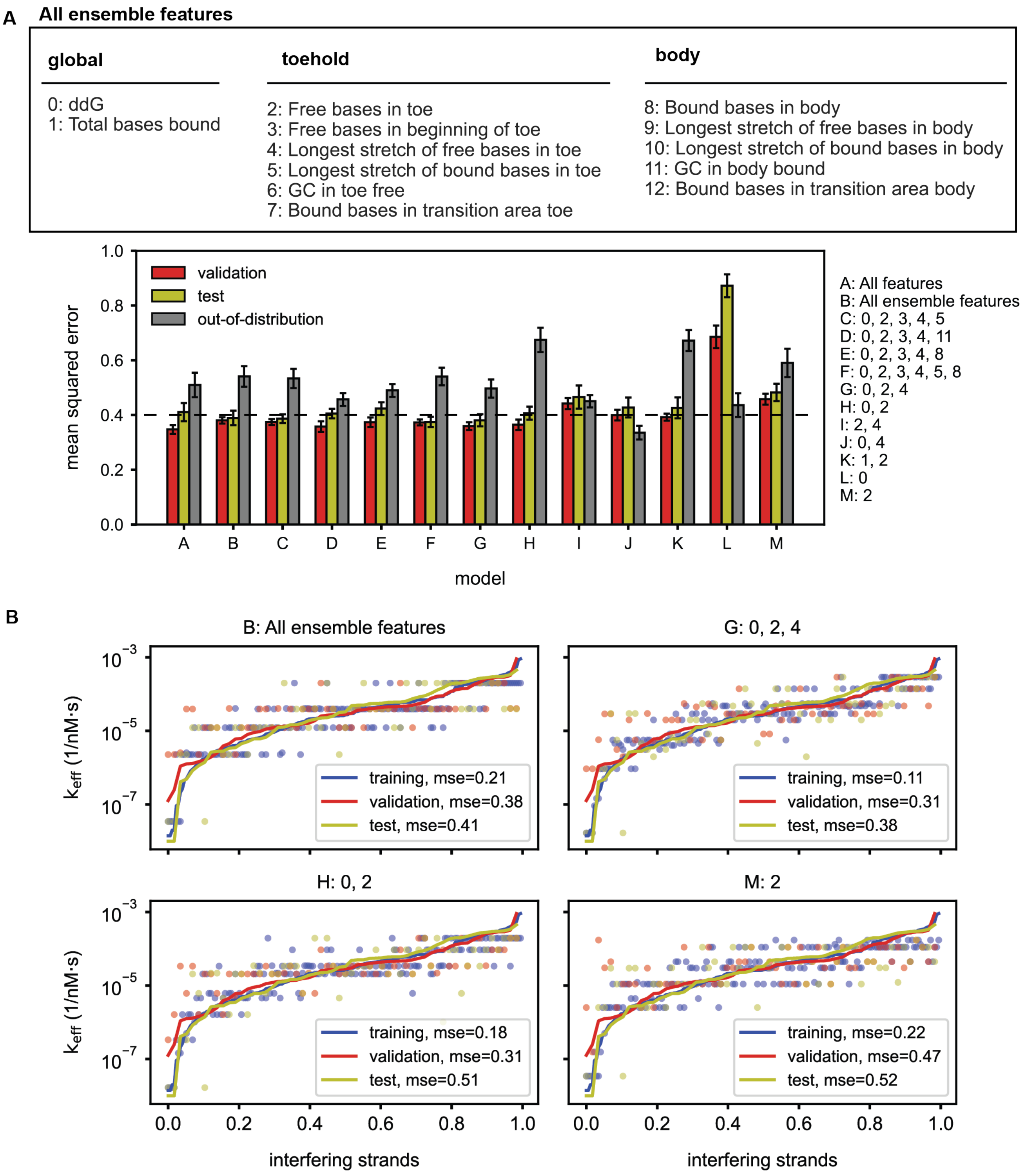

**Figure S7. Regression trees for predicting kinetics. A)** Performance of regression trees for different choices of input features trained on pooled data from two measured circuits. The shown value is the mean across ten different splits of the data into training, validation and test sets. Performance was evaluated on the validation and test set as well as out-of-distribution strands derived from the 1% mimic pool. The error bars are the standard deviation of the mean. The dashed line at mse=0.4 serves as a guide to the eye. **B)** Predictions for the best-performing trees (as measured on the validation set) for a specific data split and choice of input features. solid line=true value, dots=predicted value, mse=mean squared error

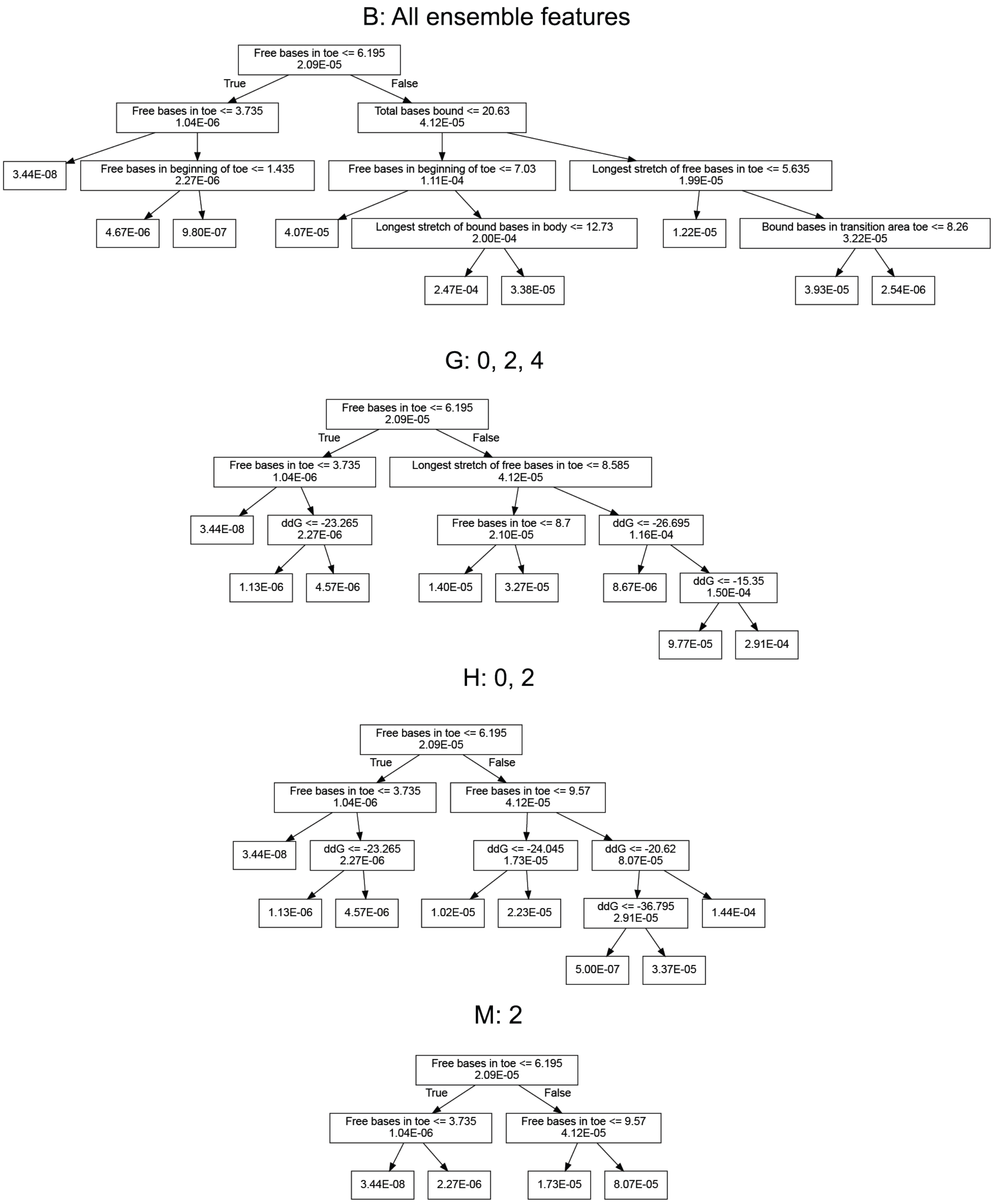

**Figure S8. A sample of specific regression trees.** The best-performing decision trees (as evaluated on the validation set) for a specific data split and choice of input features are shown.

A

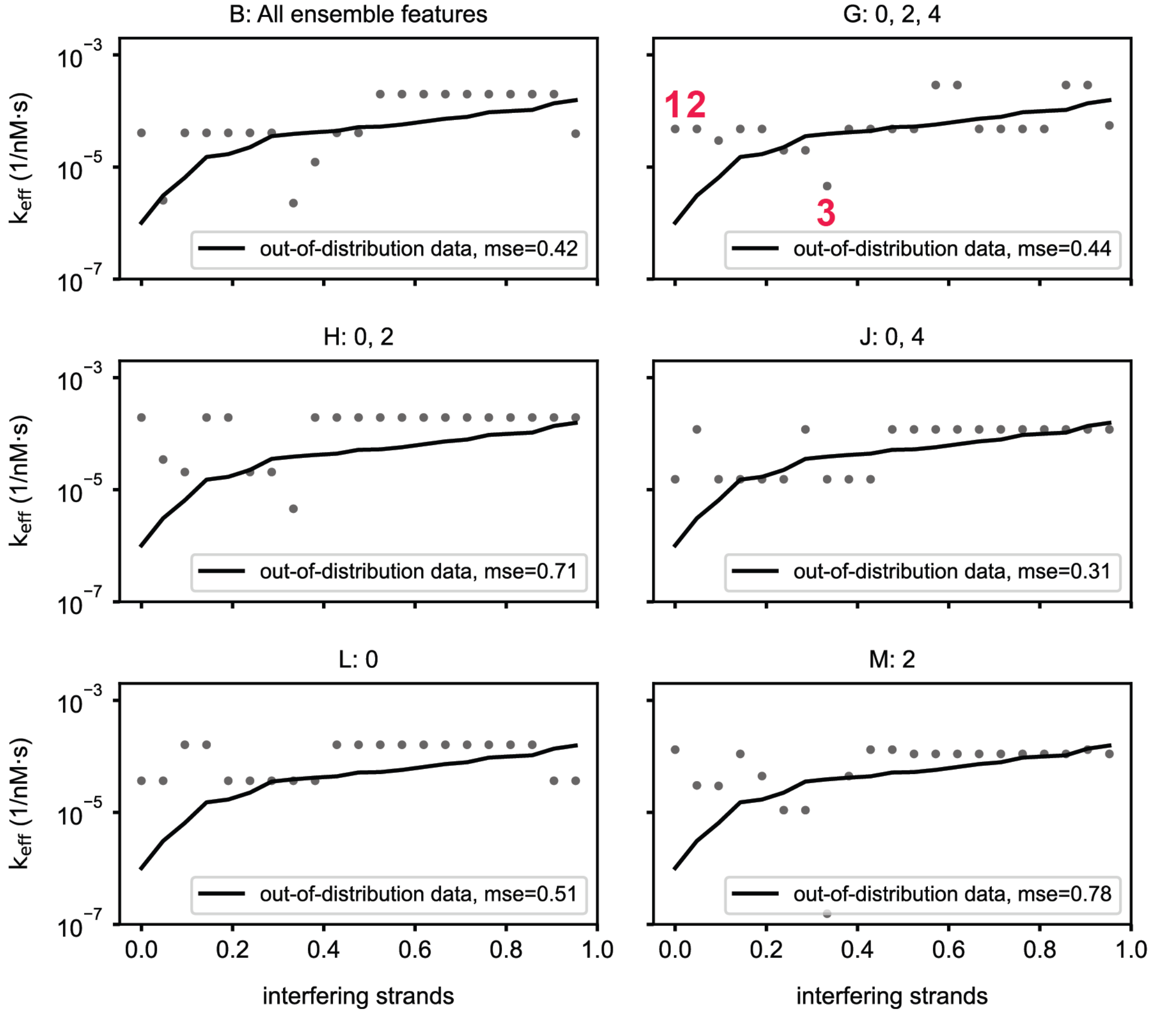

B

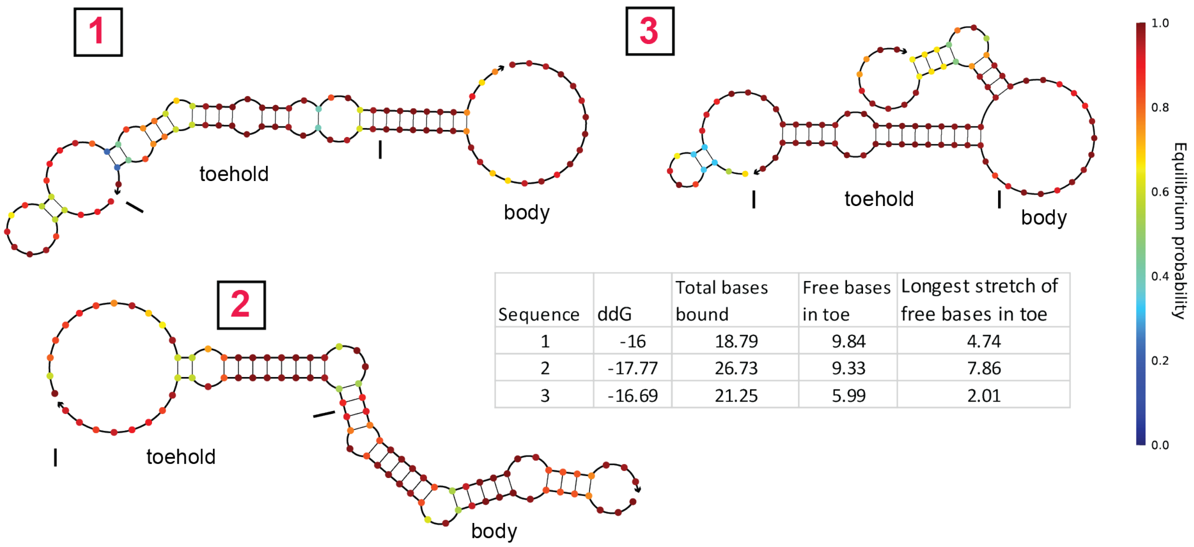

| Sequence | ddG | Total bases bound | Free bases in toe | Longest stretch of free bases in toe |
|---|---|---|---|---|
| 1 | -16 | 18.79 | 9.84 | 4.74 |
| 2 | -17.77 | 26.73 | 9.33 | 7.86 |
| 3 | -16.69 | 21.25 | 5.99 | 2.01 |

**Figure S9. Regression tree performance for out-of-distribution data. A)** Specific predictions for the best-performing trees (as measured on the validation set) for a specific data split in the training set and choice of input features applied to out-of-distribution data (i.e., strands derived from the 1% mimic pool). solid line=true value, dots=predicted value, mse=mean squared error **B)** Failure analysis for specific strands (predictions are marked for model G in A)). The bottom strand is the 45 nt long invader, the top strand the 50 nt long interfering strand.

**All ensemble features**

| global | toehold | body |
|---|---|---|
| 0: ddG | 2: Free bases in toe | 8: Bound bases in body |
| 1: Total bases bound | 3: Free bases in beginning of toe | 9: Longest stretch of free bases in body |
| | 4: Longest stretch of free bases in toe | 10: Longest stretch of bound bases in body |
| | 5: Longest stretch of bound bases in toe | 11: GC in body bound |
| | 6: GC in toe free | 12: Bound bases in transition area body |
| | 7: Bound bases in transition area toe | |

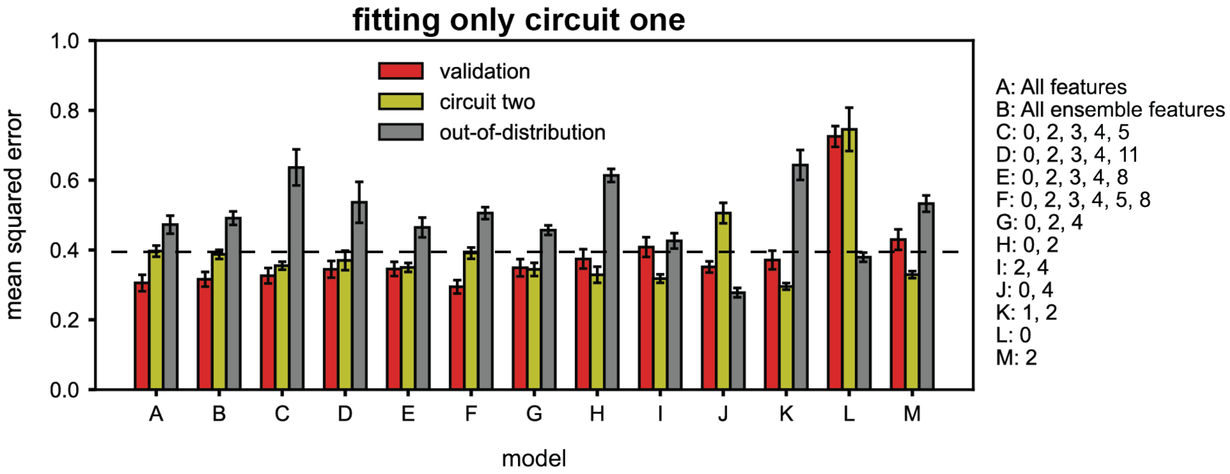

**Figure S10. Regression tree performance for split circuit one and circuit two data.** The decision trees for different choices of input features were trained on all data from circuit one. The shown value is the mean across ten different splits of the data into training and validation sets. Performance was evaluated on the validation set, all data from circuit two, as well as out-of-distribution strands derived from the 1% mimic pool. The error bars are the standard deviation of the mean. The dashed line at mse=0.4 serves as a guide to the eye.

### Symbolic Regression

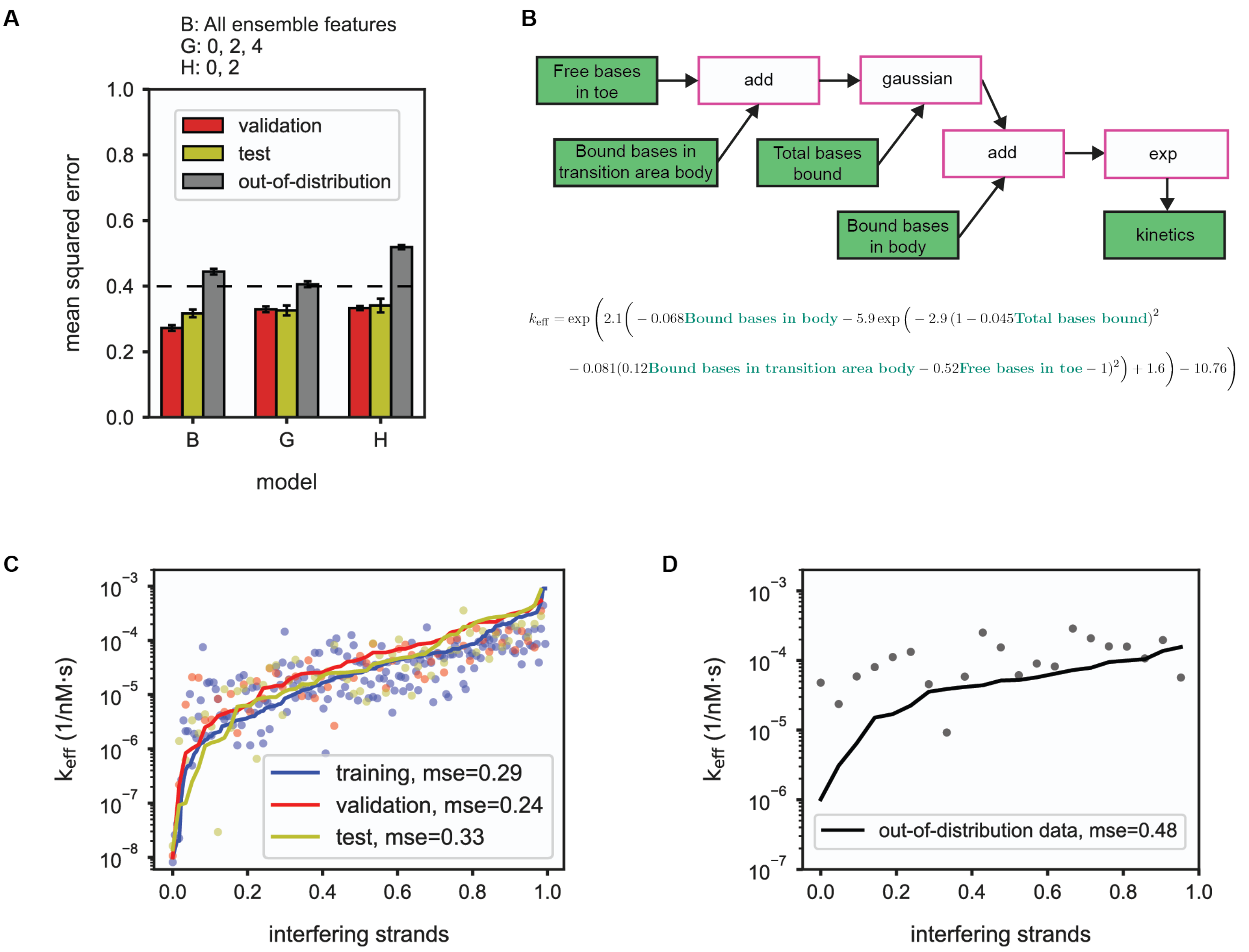

**Figure S11. Symbolic regression using Feyn. A)** Performance for different choices of input features trained on pooled data from two measured circuits. The shown value is the mean across ten different splits of the data into training, validation and test sets. Performance was evaluated on the validation and test set as well as out-of-distribution strands derived from the 1% mimic pool. The error bars are the standard deviation of the mean. The dashed line at mse=0.4 serves as a guide to the eye. **B)** Diagram and equivalent formula of a specific model for the full set of ensemble features. **C, D)** Predictions for a specific data split of the formula shown in B) for C) data from circuit one and two and D) out-of-distribution strands derived from the 1% mimic pool. solid line=true values, dots=predicted values, mse=mean squared error

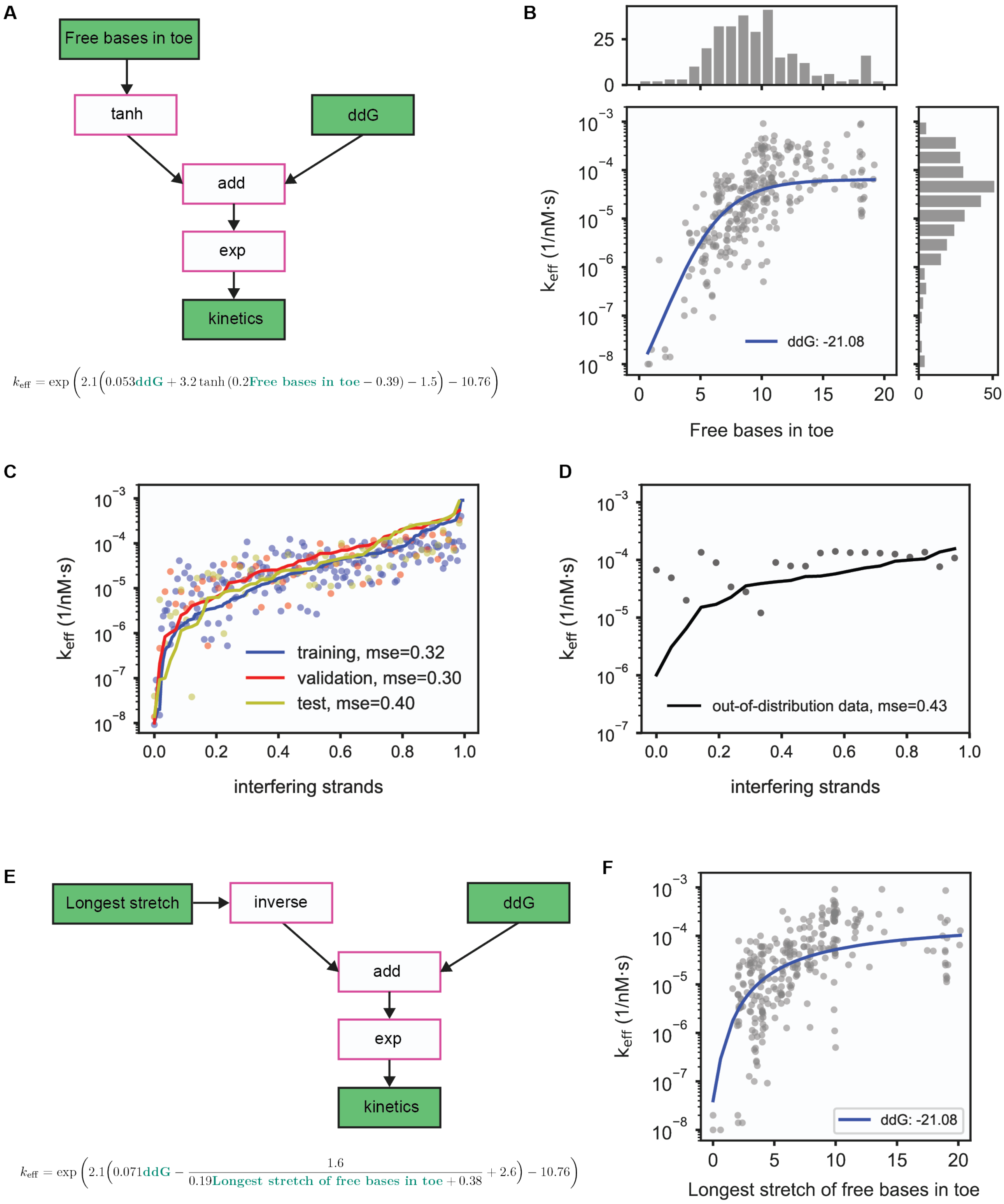

$$k_{\mathrm{eff}} = \exp\left(2.1\left(0.053\mathbf{ddG} + 3.2\tanh\left(0.2\mathbf{Free\ bases\ in\ toe} - 0.39\right) - 1.5\right) - 10.76\right)$$

$$k_{\mathrm{eff}} = \exp\left(2.1\left(0.071\mathbf{ddG} - \frac{1.6}{0.19\mathbf{Longest\ stretch\ of\ free\ bases\ in\ toe} + 0.38} + 2.6\right) - 10.76\right)$$

**Figure S12. A simple formula for estimating kinetics. A)** Diagram and equivalent formula of a simple kinetic model derived by Feyn. **B)** Model prediction for a different number of free bases in the toehold with the ddG value fixed at the median of the data. solid line=predicted values, dots=true values **C, D)** Predictions for a specific data split of the formula shown in A) for C) data from circuit one and two and D) out-of-distribution strands derived from the 1% mimic pool. solid line=true values, dots=predicted values, mse=mean squared error **E)** Diagram and equivalent formula of a simple kinetic model derived by Feyn. **F)** Model prediction for different longest stretches of free bases in the toehold with the ddG value fixed at the median of the data. solid line=predicted values, dots=true values. Note that (in contrast to the other Figures), in B) and F) the prediction is shown as a solid line to match the representation in Ref. 2 for better comparison.

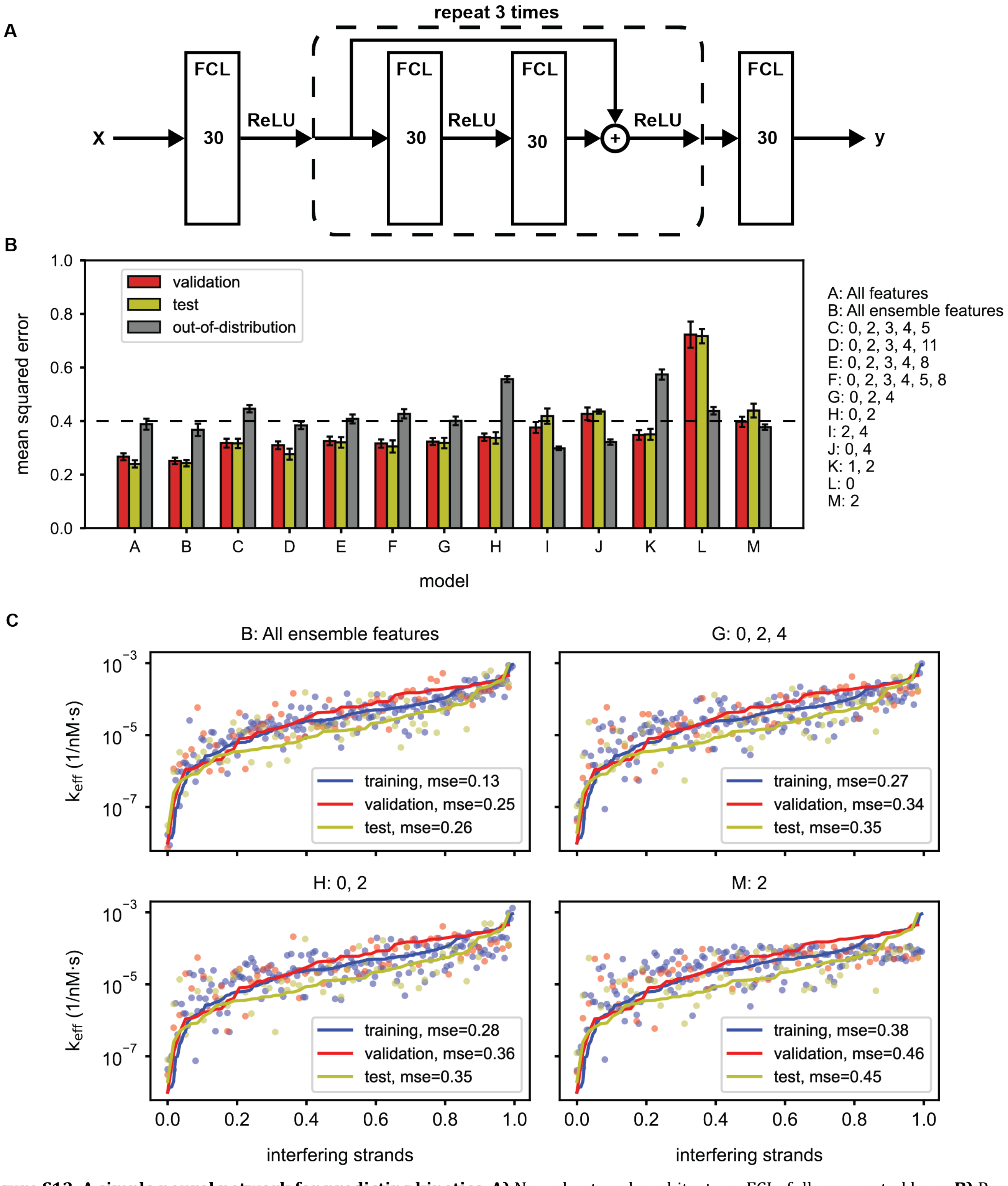

**Figure S13. A simple neural network for predicting kinetics. A)** Neural network architecture. FCL=fully connected layer **B)** Performance for different choices of input features trained on pooled data from two measured circuits. The shown value is the mean across ten different splits of the data into training, validation and test sets. Performance was evaluated on the validation and test set as well as out-of-distribution strands derived from the 1% mimic pool. The error bars are the standard deviation of the mean. The dashed line at mse=0.4 serves as a guide to the eye. **C)** Predictions for a specific data split and choice of input features. solid line=true values, dots=predicted values, mse=mean squared error

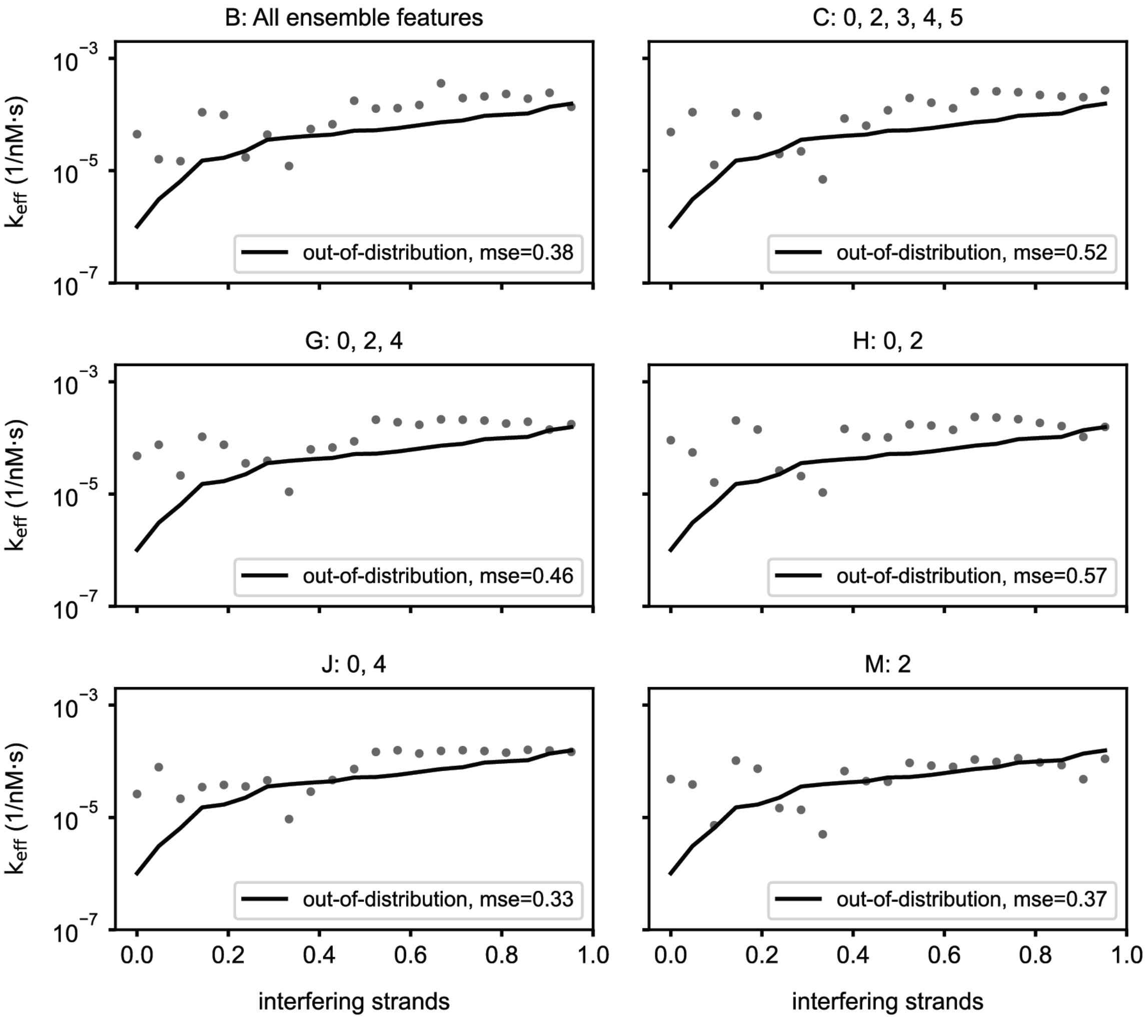

**Figure S14. Neural network performance for out-of-distribution data.** The neural network was trained for different choices of input features on pooled data from two measured circuits and evaluated on out-of-distribution strands derived from the 1% mimic pool. solid line=true values, dots=predicted values, mse=mean squared error.

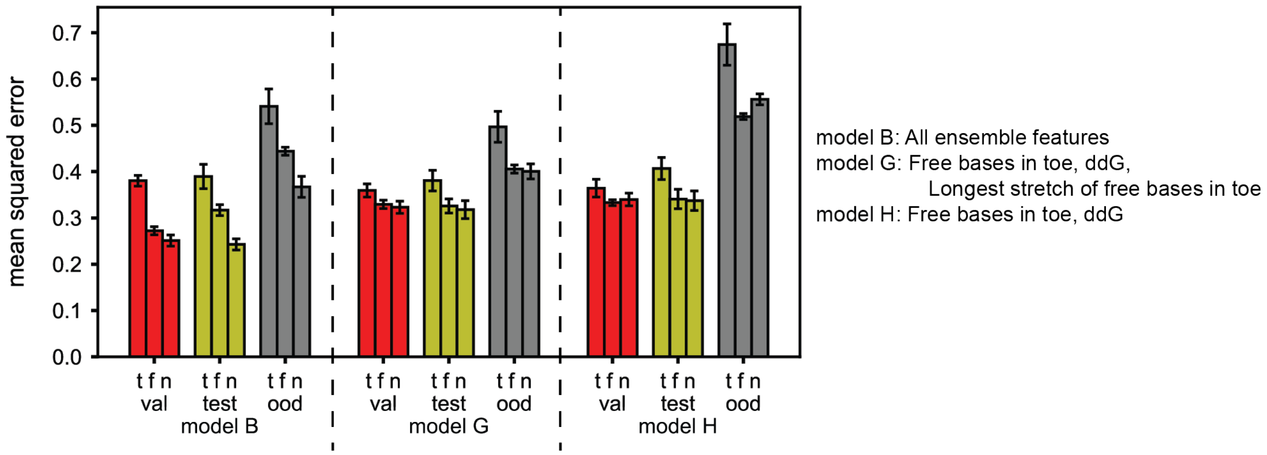

**Figure S15. Comparison between prediction models.** The prediction error on validation (val), test, and out-of-distribution (ood) data sets is evaluated for three different choices of input features (models). t = regression tree, f = Feyn, n = neural network

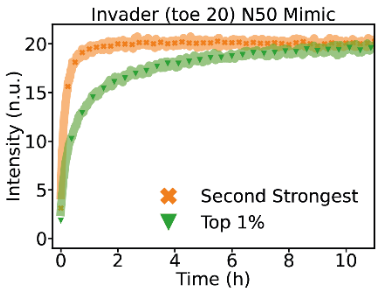

**Figure S16. Comparison between using only and all but the top 1% binders for kinetics measurements.** Green: Kinetic curve using only the upper 1% of the ΔΔG distribution. Orange: Using strands from the 8% being part of the second strongest ΔΔG regime (Figure 4). The green curve shows much slower kinetics meaning, that the top 1% have a much greater impact on displacement kinetics.

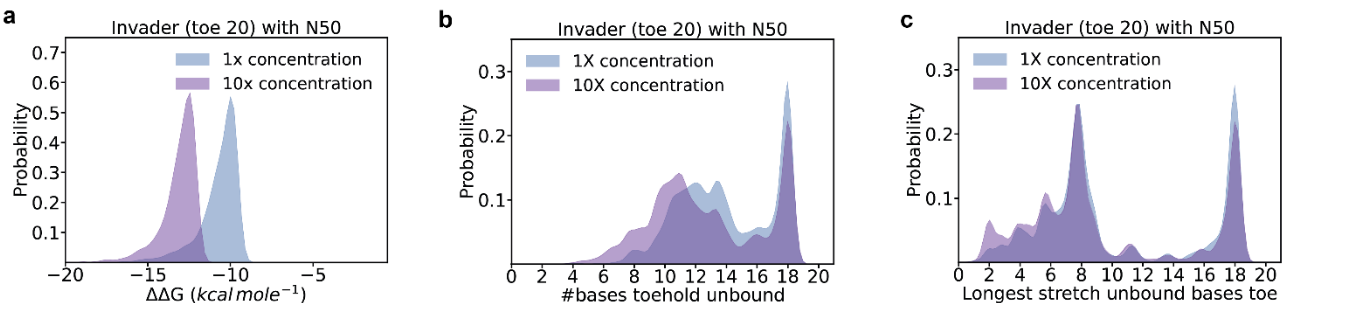

**Figure S17. Feature extraction for an invader (toe 20) interacting with N50 at different pool concentrations.** The concentration was altered by using 10 times as many strands for NUPACK calculations (10000 for 1x and 100000 for 10x). This is valid as the number of possible different strands for N50 is much higher than the number of strands physically present in the µM concentration range. Therefore, a higher concentration leads to a larger number of different strands instead of multiple copies of the same strand. We then took the top 1000 Strands and extracted **a)** the ΔΔG values, **b)** the number of unbound bases in the toehold domain, **c)** the longest stretch of unbound bases in the toehold domain. Overall, a higher concentration leads to more complexes with lower ΔΔG values and more bases bound in the toehold domain. However, binding of all toehold bases is still very unlikely as 20 consecutive bases have to be perfectly complementary. Therefore, the reaction is still expected to reach completion with every invader strand eventually bound to a substrate strand.

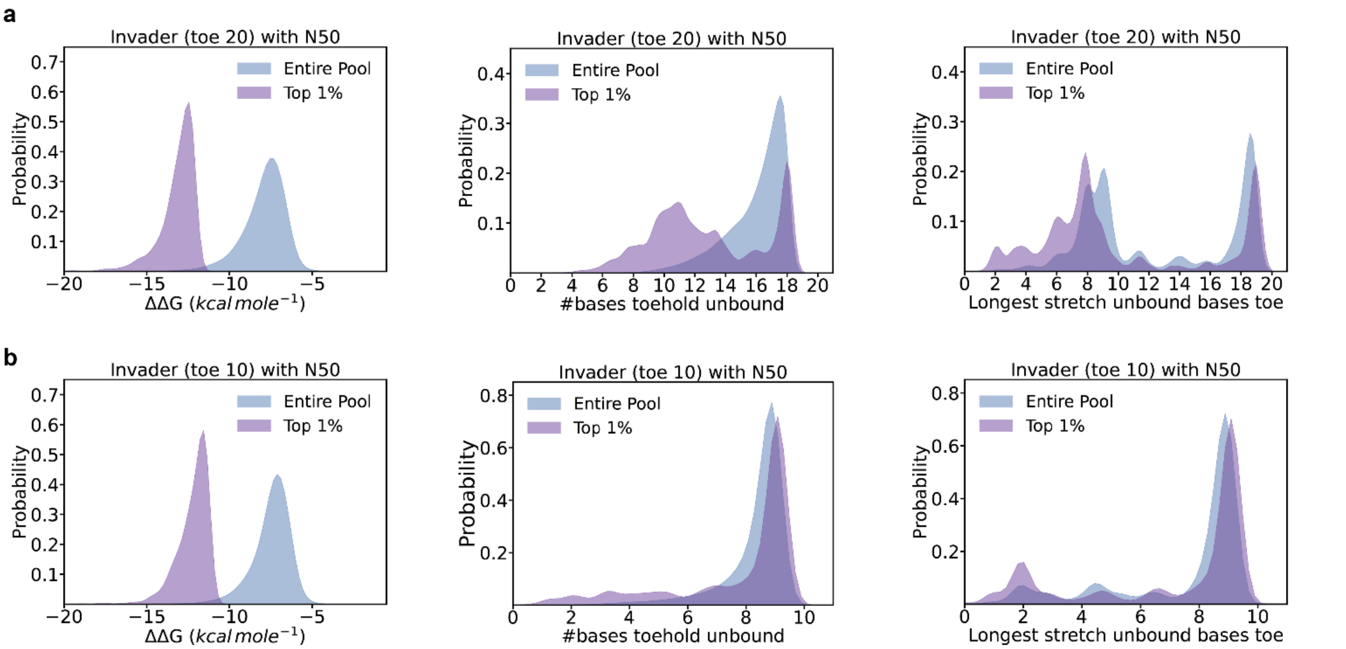

**Figure S18. Comparison of different features for an entire random pool and the top 1% of binding strands.** Three features (ΔΔG, free bases in toehold, and longest stretch of free bases in toehold) are evaluated for invaders with toehold lengths of **a)** 20 and **b)** 10. For both toehold lengths, there is a clear shift towards fewer free bases and shorter stretches of unbound bases in the toehold domain for the top 1% of strands.

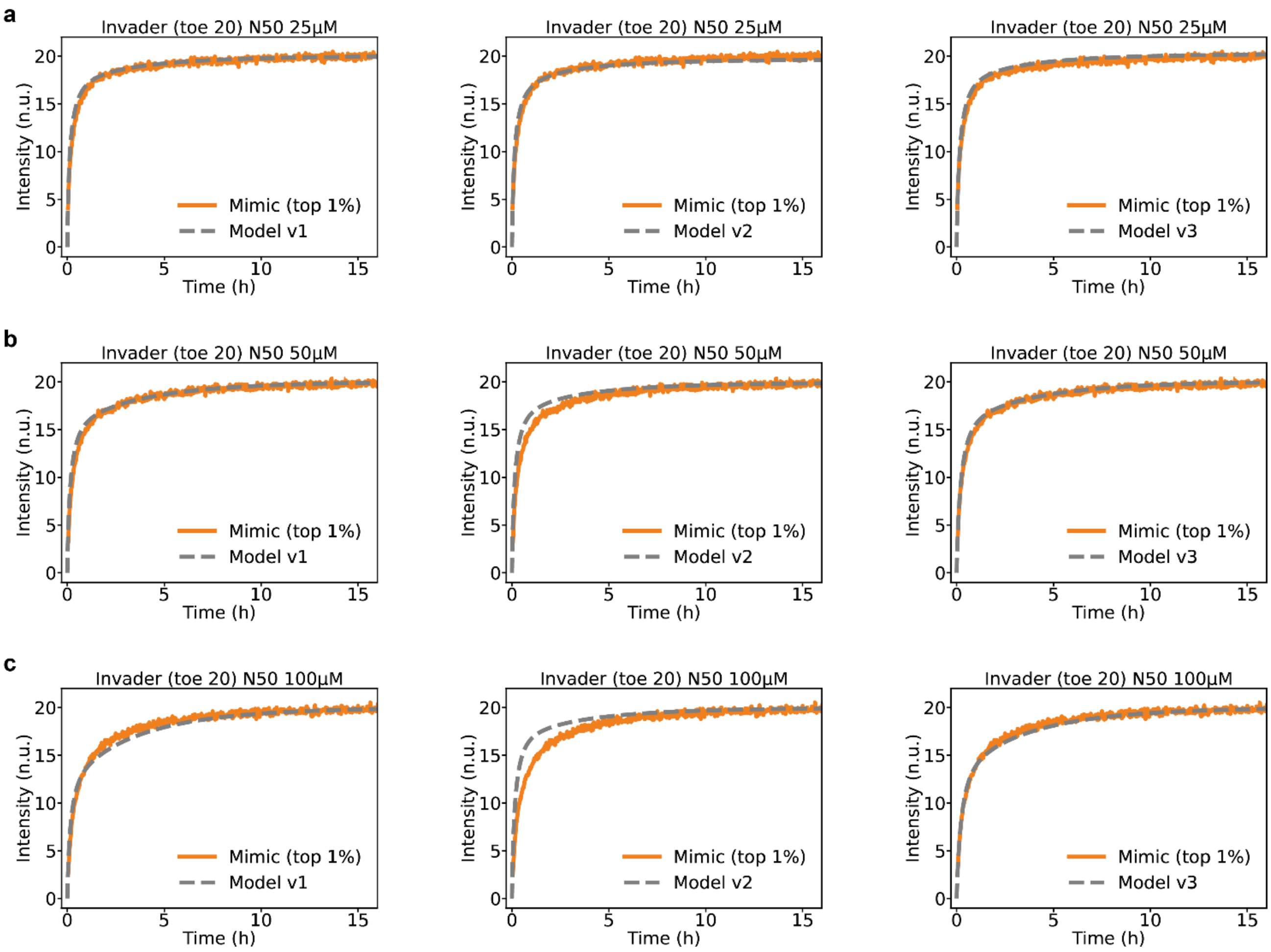

**Figure S19. Comparison of the times to reach t0.8 for measurements of the mimic pool and ODE simulations for different model versions.** As the initial concentrations are highly sensitive to small changes in the ΔΔG values, we tested two different energy models for NUPACK and also allowing complexes consisting of more than two strands. The two energy models are the default model with coaxial stacking and no dangle treatment (version 1) and a second model with "some" dangle and no coaxial stacking (version 2). We saw a better fit to our data with the version 1 having no dangle treatment. We repeated the analysis with energy model [1], but allowing for complexes with up to three interacting strands, which revealed that two strands of our strongest 20 almost always bind an invader in such a three-strand complex (version 3). We additionally determined the ΔΔG value and measured the $k_{eff}$ value for this complex in an individual measurement and used this value in the model, which slightly increased the accuracy. As the description appeared to be the physically most accurate, we used model version 3 in the main text. Subfigures **a-c)** show the kinetic curves for the mimic (top 1%) and the three different model versions for the concentrations of 25, 50 and 100µM respectively.

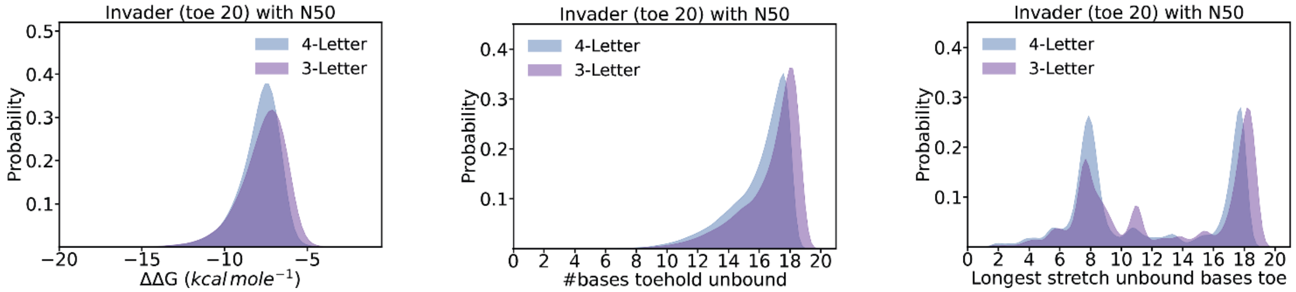

**Figure S20. Comparison of different features for a 3- and a 4-Letter alphabet invader.** Three features ($\Delta\Delta G$, free bases in toehold, and longest stretch of free bases in toehold) are evaluated for invaders with a regular 4-Letter alphabet and a 3-Letter alphabet without Cs. The sequences were chosen such that every C in the 4-Letter alphabet invader can be replaced by a G without introducing secondary structure, keeping the overall GC content the same and both strands secondary structure-free while making the sequences otherwise highly similar and, therefore, comparable. The interaction of the 3-Letter invader with the random sequences is slightly less stable (higher ΔΔG values) and the toehold domain slightly more accessible, which indicates that the 3-Letter alphabet invader should display faster TMSD kinetics in random pools.
